# Suppression of pyramidal neuron G protein-gated inwardly rectifying K+ channel signaling impairs prelimbic cortical function and underlies stress-induced deficits in cognitive flexibility

**DOI:** 10.1101/2020.06.08.139725

**Authors:** Eden M Anderson, Steven Loke, Benjamin Wrucke, Annabel Engelhardt, Evan Hess, Kevin Wickman, Matthew C Hearing

## Abstract

**Background:** Imbalance in prefrontal cortical (PFC) pyramidal neuron excitation:inhibition is thought to underlie symptomologies shared across stress-related disorders and neuropsychiatric disease, including dysregulation of emotion and cognitive function. G protein-gated inwardly rectifying K^+^ (GIRK/Kir3) channels mediate excitability of medial PFC pyramidal neurons, however the functional role of these channels in mPFC-dependent regulation of affect, cognition, and cortical dynamics is unknown.

**Methods:** In mice harboring a ‘floxed’ version of the kcnj3 (Girk1) gene, we used a viral-cre approach to disrupt GIRK1-containing channel expression in pyramidal neurons within the prelimbic (PL) or infralimbic (IL) cortices. Additional studies used a novel model of chronic unpredictable stress (CUS) to determine the impact on PL GIRK-dependent signaling and cognitive function.

**Results:** In males, loss of pyramidal GIRK-dependent signaling in the PL, but not IL, differentially impacted measures of affect and motivation, and impaired working memory and cognitive flexibility. CUS produced similar deficits in affect and cognition that paralleled a reduction in PL pyramidal GIRK-dependent signaling akin to viral approaches. Viral- and stress-induced behavioral deficits were rescued by systemic injection of a novel, GIRK1-selective agonist, ML-297. Unexpectedly, neither ablation of PL GIRK-dependent signaling or exposure to the CUS regimen impacted affect or cognition in female mice.

**Conclusions:** GIRK-dependent signaling in male mice, but not females, is critical for maintaining optimal PL function and behavioral control. Disruption of this inhibition may underlie stress-related dysfunction of the PL and represent a therapeutic target for treating stress-induced deficits in affect regulation and impaired cognition that reduce quality of life.

## INTRODUCTION

Dysfunction of the prefrontal cortex (PFC) is an underlying factor in both affect and cognition-related behavioral deficits that co-occur across neuropsychiatric disorders (1–12). Similar symptomologies are observed in individuals with chronic psychosocial or self-perceived chronic stress (13–16). Converging evidence indicates that prolonged exposure to environmental stressors can increase the risk and severity of these neuropsychiatric disorders -- highlighting a need to identify neural substrates that contribute to optimal PFC function and behavioral control and how disruption in these mechanisms contributes to pathological states.

The prelimbic cortical subdivision (PL) of the medial prefrontal cortex (mPFC) is involved in top-down regulation of behavior related to anxiety, motivation, stress, and coordination of working memory and behavioral flexibility (e.g., strategy shifting) (17–26). Optimal function of the PL relies on coordinated activity of principle glutamatergic pyramidal neurons. This activity is critically dependent on a dynamic balance of cell excitation:inhibition mediated largely by intrinsic (physiological) membrane properties that influence the cellular response to synaptic input (27–31). Accordingly, imbalances in pyramidal neuron excitation:inhibition are thought to contribute to numerous symptoms observed in neuropsychiatric disorders (4, 28, 32–40).

PL pyramidal excitability and spike firing is modulated by activity of G protein-gated inwardly rectifying K^+^ (GIRK/Kir3) channels which produce a slow hyperpolarizing current that acts as a neuronal “off switch” (41–44). GIRK channels are the primary postsynaptic downstream effector for inhibitory metabotropic receptors, including perisomatic GABA_B_ receptor (GABA_B_) – a major target of local GABA neurons (45, 46). Converging clinical and preclinical data highlights a link between hypofunction of GABA_B_-GIRK signaling with cognitive disabilities, abnormalities in affect, and susceptibility to neuropsychiatric disease (44, 47–56), however the anatomical locus of this hypofunction remains unclear. As the role of GIRK signaling in PFC-dependent behavior has not been characterized, this study focused on testing the hypothesis that PL GIRK-dependent signaling is a critical locus of affect and cognitive flexibility. Further, we hypothesized that chronic unpredictable stress (CUS) would result in reductions in PL GIRK-dependent signaling and therefore deficits cognitive flexibility.

## METHODS

### Animals

All animal experimentation was approved by the Institutional Animal Care and Use Committee at Marquette University. Girk1^flox/flox^ stock mice were generated as described (57, 58), with experimental mice bred in house (91 ± 10 PD at start of ASST). For stress-related studies, adult male and female mice (91 ± 15 PD at start of ASST) were a combination of C57BL/6J bred in house, positive for a fluorescent reporter gene but with a C57BL/6J background or purchased directly from Jackson Laboratories. Mice were housed in a temperature and humidity-controlled room with a 12h/12h light/dark cycle with food and water available *ad libitum* except where described, and all procedures were conducted during the light phase. For studies requiring food deprivation, mice were maintained at 85-90% free feeding weight.

### Viral ablation of GIRK signaling

Adult male and female Girk^flox/flox^ mice were anesthetized with isoflurane and received a bilateral infusion of either AAV8:CamkII:Cre (UNC) or AAV8:CamkII:GFP (UNC) into either the PL (AP: +1.80, ML:±0.40, DV:-2.30) or IL (AP:+1.80, ML:±0.40, DV:-3.20) using a 5μL flat Hamilton syringe (0.5μL/infusion;0.1μL/min). Syringes were left in place for 5min to reduce backflow then drawn to the surface over 5min. Mice recovered for a period of at least four weeks to ensure full viral expression.

### Behavioral Testing

#### Elevated Plus Maze (EPM) and Forced Swim Test (FST)

Tests were conducted as previously described (59).

#### Forced Alteration T-Maze Paradigm

A subset of cre- and cre+ mice underwent 3d of habituation and training prior to testing in a T-maze based on previous literature (60). Following food restriction and context habituation, mice were allowed to explore the maze and consume a reward (50% diluted liquid vanilla Ensure^®^) in each arm for 5min (x5 trials). On day three, mice underwent forced run training with only one arm baited and the other blocked for 12 trials (6/arm). During the testing period, animals underwent 12 daily trials with access to one reward arm and allowed to consume the reward. Following a 20sec period in the start box, mice were given access to both arms with the previously unentered arm baited (correct choice).

#### Attention Set Shifting

Operant attentional set-shifting task (ASST) procedures were performed in sound attenuating boxes (Med Associates, Inc.) based on previously developed rat protocols (26, 61). Mice were food deprived and underwent initial food training where the left and/or right lever required pressing a combined 50 times within a 3hr and subsequent 30min session. During food training, all lever presses resulted in 20sec access to a liquid dipper containing Ensure^®^. *Lever training* was conducted with the left or right lever presented in a pseudorandom order for a total of 90 trials (45 trials/lever). Response on the presented lever resulted in access to Ensure^®^ for 20sec and no response within 10sec counted as an omission (criterion is ≤ 5 omissions on 2 consecutive days). Lever preference was assessed in a *bias test* where both the left and right lever was presented and reinforced for a total of 7 trials.

Test days for the ASST involved visual cue, extradimensional shift (ED Shift), and reversal test on separate days. Criteria for each test was 10 consecutive correct responses in a row with a minimum of 25 trials (or until 150 trials were conducted). If criterion was not reached in 150 trials, the test was conducted again the following day (5d maximum). During the *visual cue test*, a light was illuminated above the right or the left lever for 3sec followed immediately by presentation of the lever, with the correct response resulting in a reward. In the *ED Shift* and *response testing*, the cue was presented but the correct response was the lever opposite the bias regardless of cue location. In a subset of mice, 20mg/kg ML297 was injected 30min prior to the ED shift. During *reversal testing*, reinforced lever was opposite to the correct lever during the ED Shift.

#### Progressive Ratio

Following the conclusion of ASST animals remained at 85-90% free feeding weight. Responses on the left lever were reinforced and the responses required to obtain the next reward progressively increased with each reward obtained ((5e^.2*n^)-5; (62)). Testing continued until 30min elapsed without a response or a total of 1.5hr.

### Slice Electrophysiology

Mice were anesthetized with isoflurane and brains sliced in oxygenated sucrose ACSF slush as described (59). All recordings were performed using borosilicate electrodes (2.5-4.5 MΩ) filled with a K-Gluconate solution (41, 59). For rheobase and spike frequency, a 20pA current-step injection was used (0-400pA). For ML297 and baclofen recordings, a consistent holding current was obtained (<20% fluctuation), followed by bath application of 10μM ML297 in 0.04% DMSO or 200μM baclofen (Sigma-Aldrich). Evoked currents were reversed using bath application of 0.30mM barium chloride (Fisher Scientific). Recordings were filtered at 2kHz and sampled at 20kHz, with series resistance (<40MΩ) monitored throughout all recordings.

### Assessment of Viral Expression

Only mice with bilateral viral expression primarily confined to the PL or IL were used in analysis which was either confirmed in slices immediately prior to slice electrophysiology studies or in 100μm fixed tissue (4% paraformaldehyde for 24-48 hours).

### Chronic Unpredictable Stress

Male and female C57BL/6J mice received four weeks of CUS or were handled twice daily (controls), as previously described (59). Mice began training in ASST 3-5d following the last stress exposure. Following completion of testing (22-25 days post stress), a subset of mice were euthanized for slice electrophysiology.

### Data Analysis

Data are presented as mean ± SEM. Statistical analyses were performed using SigmaPlot 11.0. Independent-samples t-tests, one-way ANOVA, or repeated-measures were used; when parametric normality tests were violated, a Kruskal-Wallis or Mann-Whitney nonparametric test was used. Student-Newman-Keuls method for multiple post-hoc comparisons was used when applicable. Statistical outliers (±2SD) were excluded from analyses (11 data points from all experiments). Electrophysiology sample sizes are denoted as *n* for the number of recordings/cells and *N* for the number of mice.

## RESULTS

### Characterization of PL Pyramidal Neuron GIRK Knockout

GIRK channels containing the GIRK1 subunit mediate a majority of GIRK-dependent signaling in layer 5/6 (L5/6) PL pyramidal neurons and loss of this signaling leads to enhanced excitability (41). To validate our model, we began by evaluating GABA_B_- and GIRK1-dependent signaling in L5/6 pyramidal neurons in GIRK1^flox/flox^ male mice bilaterally infused with a cre- expressing or GFP-expressing adeno-associated virus (AAV) driven by the CaMKII promoter (Figure 1a). In GFP-expressing cells (cre-), bath application of the GABA_B_ agonist, baclofen (200μM), evoked an outward current (I_Baclofen_) that was reversed by a low concentration of external Ba^2+^ (0.30mM), while fluorescently identified pyramidal neurons from Girk1^flox/flox^ mice expressing cre (cre+) had a significant reduction in I_Baclofen_ mean amplitude (Figure 1b). Similarly, GIRK currents evoked by bath application of the GIRK1-selective agonist, ML297, were significantly reduced (~84%) in cre+ pyramidal neurons compared to cre-pyramidal neurons (Figure 1c). In a subset of cells, bath application of DMSO vehicle alone produced a current similar in amplitude to residual I_ML297_ in cre+ PYR (n=3, 37.7 ± 5.8 pA; data not shown), suggesting that the remaining current is largely vehicle-dependent.

**Figure 1.**
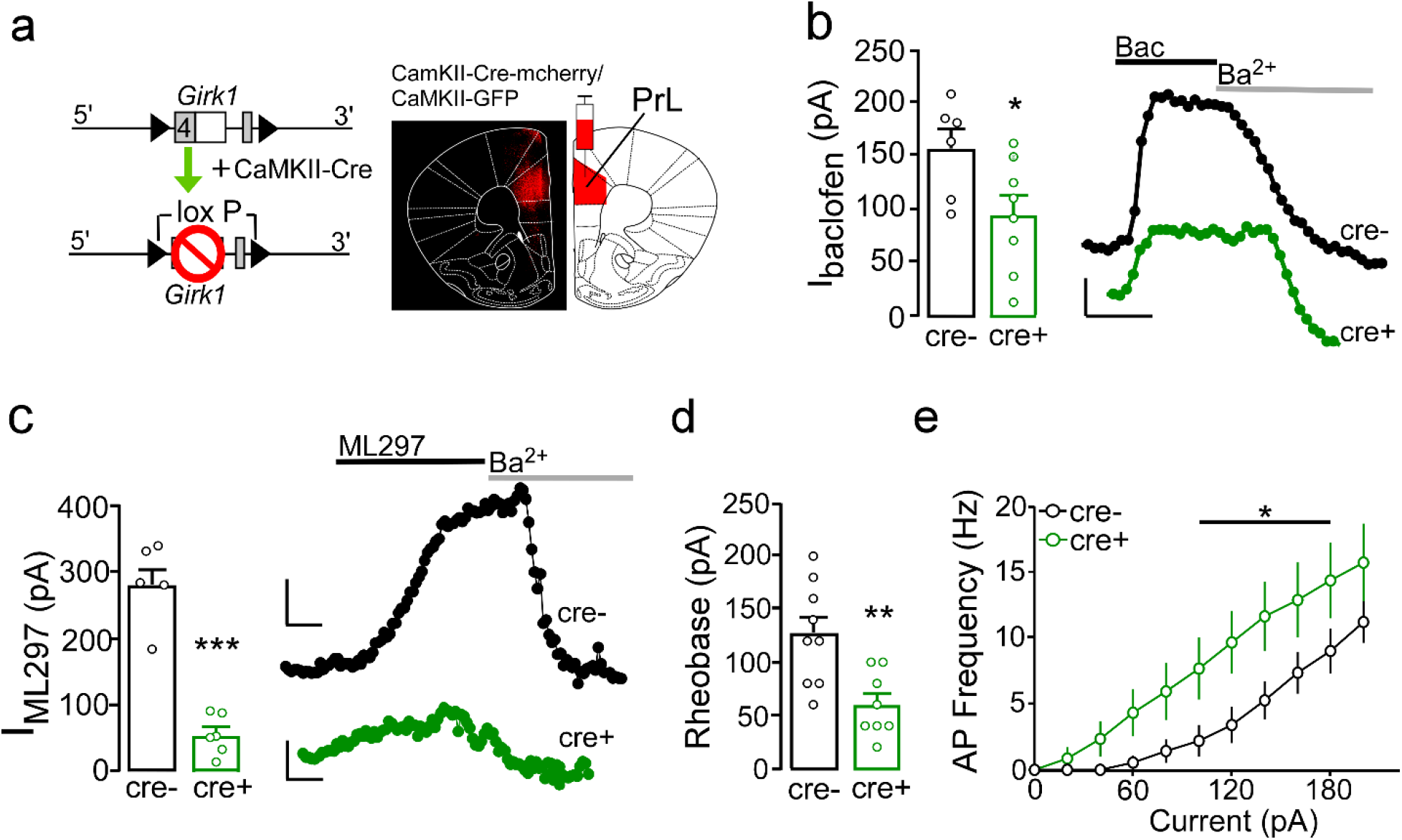
**a.** Schematic showing knockdown of GIRK1 in the PrL PYR through bilateral infusion of cre+ or gfp-expressing AAV driven by the CaMKII promotor with L5/6 pyramidal neurons recorded from using whole-cell slice electrophysiology. **b.** Outward current evoked by bath application of baclofen (I_baclofen_; 200μM) in Girk1^flox/flox^ male mice were significantly reduced in cre+ PrL L5/6 PYR (t_(12)_= 2.33, p=0.038; n=8/N=5) versus cre- (n=7/N=4). I_baclofen_ was reversed in the presence of external Ba^2+^ (0.30mM). **c.** Outward current evoked by external application of the GIRK1 agonist, ML297 (I_ML297_) was reduced in cre+ Girk1^flox/flox^ L5/6 PL pyramidal neurons (t_(10)_= 8.67, p<0.001; n=6/N=4) compared to cre- (n=6/N=5). **d.** Threshold to fire an action potential (rheobase) in PL L5/6 pyramidal neurons was reduced in cre+ (t_(16)_= 3.63, p=0.002; n=8/N=6) compared to cre- (n=10/N=7). **e.** Current-spike plots from Girk1^flox/flox^ PL L5/6 pyramidal neurons showed a condition by current interaction (F_(10,150)_= 2.23, p=0.02; cre- n=9/N=6; cre+ n=8/N=6), with cre+ pyramidal neurons exhibiting an increase in spike frequency compared to cre- at 100-180pA. *p<0.05, **p<0.01, ***p<0.001. Scale bars: 50pA x 100sec.

Having established that GABA_B_ and GIRK-dependent signaling is reduced in pyramidal neurons following viral cre treatment, we next investigated whether viral-mediated reduction in GIRK-dependent signaling in male mice is sufficient to increase intrinsic excitability. Cre+ pyramidal neurons exhibited a reduction in current required to fire an action potential (rheobase) compared to cre- pyramidal neurons (Figure 1d). Assessment of current-spike relationship showed a significant virus by current interaction (Figure 1e) with significant post-hoc comparisons at 100-180pA. Collectively, these findings verify the ability of viral-mediated approaches to suppress GIRK-dependent inhibition and increase excitability of L5/6 PL pyramidal neurons.

### PL Pyramidal Neuron GIRK Knockout variably influences performance in EPM, FST, and Progressive Ratio

Percent time spent in the open arms of an EPM, time immobile in the FST, and break point during progressive ratio were measured to determine the impact of PL pyramidal GIRK knockout in male mice (Figure 2a). Cre+ male mice spent a greater percentage of time in the open arm compared to cre- (Figure 2b). Alternatively, cre+ mice exhibited an increase in mean time spent immobile compared to cre- in the FST (Figure 2c). Using a progressive ratio test to assess motivation, we found no differences in total correct responses but a trend towards increased break point (Figure 2d). These data indicate that reducing PL pyramidal GIRK-dependent signaling in males has unique effects on affect-related behavior that reduces normal anxiety responses and escape-related strategies suggestive of an anxiolytic and pro-depressive phenotype (63, 64), while potentially increasing reward motivation.

**Figure 2.**
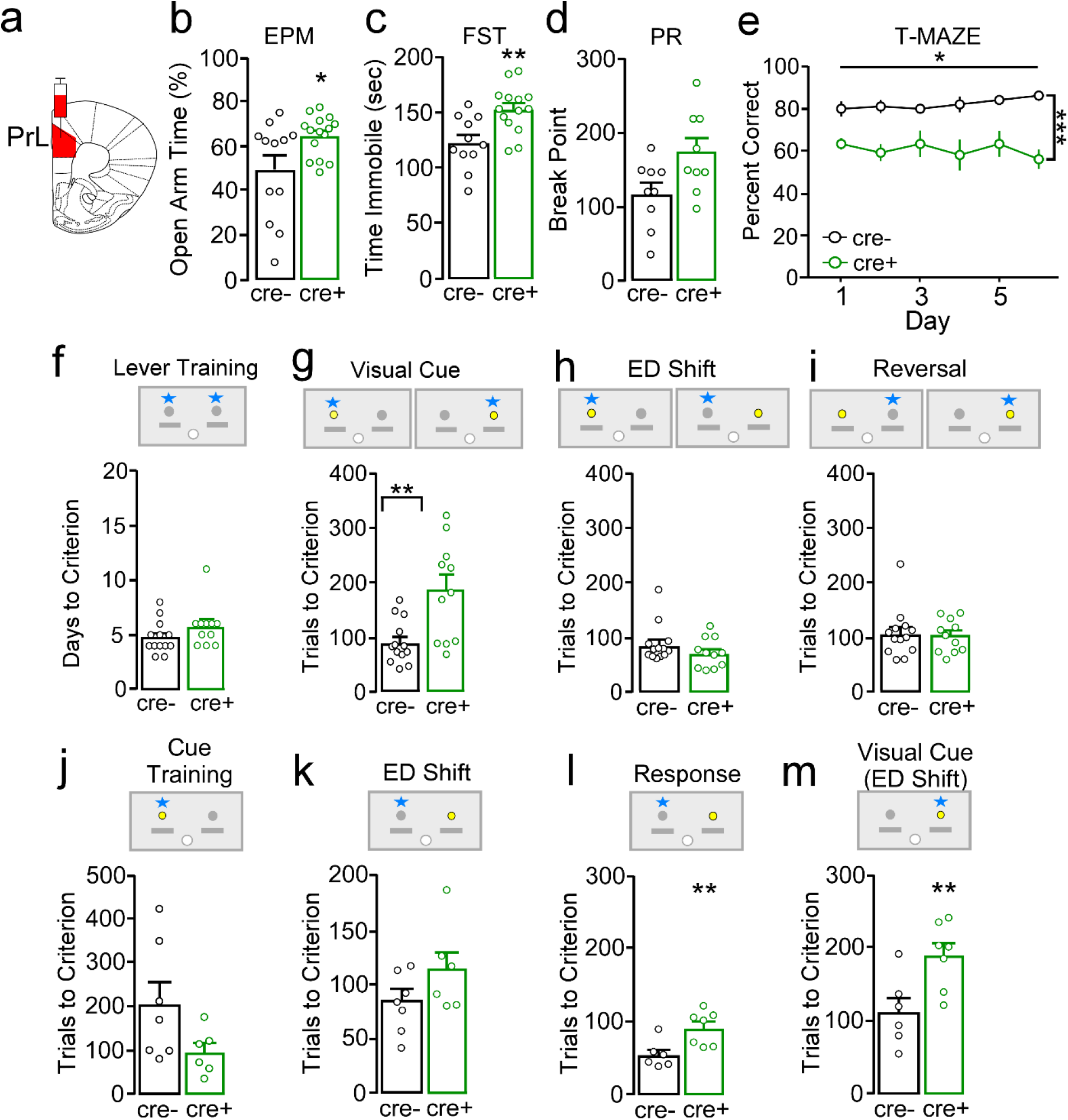
**a.** Schematic showing viral infusion into the PL. **b.** Percent time in the open arm of the EPM was increased in cre+ male mice compared to cre- controls (t_(25)_= −2.41, p=0.02). **c.** Total time (seconds) spent immobile during the FST was increased in cre+ compared to cre- mice (t_(24)_= −3.07, p=0.005). **d.** Break point for number of responses to receive liquid reward during a progressive ratio test showed a trend towards an increase in cre+ males versus cre- controls (*U*= 19.00, p= 0.06) and no difference in number of correct responses (t_(16)_= −0.98, p=0.34; data not shown). **e.** Percent correct trials (out of 12 trials) across days the forced alteration t-maze paradigm. A main effect of condition (F_(1,14)_= 26.74, p<0.001), but not day (F_(5,70)_= 0.28, p=0.92) or condition x day interaction was observed (F_(5,70)_= 0.98, p=0.43), with cre+ mice having reduced percent correct each test day. **f.** There was no difference between cre- and cre+ male mice in the number of days to reach lever training criterion (t_(22)_=-1.37, p=0.19). **g.** During the visual cue test, cre+ mice took significantly more trials to reach criterion compared to cre- mice (t_(22)_=-3.40, p=0.003). **h.** During the ED Shift test, cre- and cre+ mice took similar number of trials to reach criterion (t_(22)_=1.13, p=0.27). **i.** During the reversal test, cre- and cre+ mice took similar trials to reach criterion (t_(22)_=0.11, p=0.92). **j.** Girk1^flox/flox^ cre+ and cre- did not differ in trials to reach criterion for visual cue training (t_(11)_=1.87, p=0.09) **k.** Following visual cue training, there were also no differences in trials to reach criteria in the ED Shift (t_(11)_=-1.45, p=0.18). **l.** During lever training prior to a response then cue test, cre+ mice took longer to obtain lever training criteria compared to controls (t_(11)_= −3.28, p=0.007; data not shown). Cre+ mice required greater number of trials to reach criterion during an initial response test (t_(11)_= −3.08, p=0.01). **m.** Cre+ mice also took more trials to reach criterion during the subsequent ED response-to-cue test (t_(11)_= −2.98, p=0.01). *p<0.05, **p<0.01, ***p<0.001.

### Working memory in PL Pyramidal Neuron GIRK Knockout mice

The most consistently documented cognitive deficits in neuropsychiatric disease and stress pathologies include impaired behavioral flexibility and working memory (65, 66), thus we examined whether loss of GIRK promotes similar impairments. Using a forced alteration T-maze paradigm to assess working memory, we found no effect of day, or condition (virus) by day interaction. A main effect of condition was detected, with cre+ mice showing a reduction in percent correct choices compared to cre- controls each day (Figure 2e). These data indicate that PL pyramidal neuron GIRK1-dependent signaling in male mice plays a key role in information processing related to working memory, and that these deficits can be rescued with systemic targeting activation of GIRK1-containing channels.

### Impact of PL Pyramidal Neuron GIRK Knockout on Cognitive Flexibility

To determine if PL pyramidal GIRK channels play a role in complex forms of PFC-dependent cognition, we examined the impact of PL GIRK1 suppression on cognitive flexibility. We used a modified operant-based ASST (61) that is reliant on PL function (24, 67) and resembles the Wisconsin Card Sorting Task in its sensitivity to distinct components of decision making such as suppression of irrelevant strategies, acquisition and generation of novel strategies, and maintenance of effective strategies. To determine if GIRK1^flox/flox^ mice exhibit a baseline phenotype, we first compared GIRK1^flox/flox^ mice to C57BL/6J mice receiving sham surgery and found no difference in their performance on any ASST (lever train: t_(11)_=-0.98, p=0.35; VC trials: t_(11)_=-0.34, p=0.74; ED trials: t_(11)_=-0.93, p=0.38; REV trials: t_(11)_=-1.27, p=0.23; data not shown), thus data were combined for further analysis. There was no difference in days to criteria for lever training between cre- and cre+ (Figure 2f). Following lever training, comparison of performance in the visual cue test showed that cre+ mice required significantly greater number of trials (Figure 2g) and errors (Supplemental 1a) to reach criteria compared to cre- but did not differ on omissions (Supplemental Table 1). During the ED Shift, cre- and cre+ mice required a similar number of trials (Figure 2h), errors (Supplemental 1b) and omissions to reach criteria (Supplemental Table 1). Similarly, no difference was observed in trials (Figure 2i), errors (Supplemental 1c) or omissions (Supplemental Table 1) to criteria in a subsequent reversal learning test.

Pharmacological inhibition of the PL does not impact performance in an operant or maze-based visual cue discriminative learning task in rats (61, 68), thus to ensure that impaired performance during the visual cue test did not reflect deficits in general attention to a cue, we next trained a new set of mice using the visual cue test with similar criteria. There were no differences in the trials (Figure 2j), errors (Supplemental 2a), or omissions (Supplemental Table 2) to reach criteria between cre+ and cre- mice. After reaching criteria, mice underwent the ED Shift test where there were no differences in trials (Figure 2k), errors (Supplemental 2b), or omissions (Supplemental Table 2). There were also no differences in trials or errors to criterion (Supplemental 2c-d) or omissions to criterion (Supplemental Table 2) during the reversal test.

As mice were initially lever trained in a pseudo-random fashion and were to press either the left or the right lever when it was presented (i.e. only attend to levers), it is possible that the addition of a visual cue rule was acting as an attentional shift (i.e. ignore previous lever presentation order and attend to visual cue). To address this, additional cohorts underwent lever training during which cre+ male mice took longer to obtain criteria compared to controls (data not shown) followed by a response test (lever opposite of bias is correct) and a subsequent visual cue test. During the response test, cre+ mice took significantly more trials (Figure 2l) and errors (Supplemental 3a) however also had greater omissions (Supplemental Table 3). During the visual cue set-shift, cre+ mice also required a greater number of trials (Figure 2m) and errors (Supplemental 3b) to reach criterion compared to cre- but did not differ in omissions (Supplemental Table 3). These findings combined with a lack of difference in cue-based learning indicate that loss of GIRK channel activity alters PL function (i.e. cognitive flexibility) in a manner distinct from lesions.

### Influence of IL Pyramidal Neuron GIRK Knockout on affect and cognitive behavior

Prior research indicates that similar to the PL, the more ventral IL subdivision of the mPFC also regulates affect- and cognition-related behavior (24, 69, 70) and that imbalances in IL cortical excitation:inhibition may promote deficits in these behaviors (68, 71). Assessment of ablation of IL pyramidal neuron GIRK1-dependent signaling (Figure 3a) showed that cre+ and cre- mice spent a similar amount of time in the open arm during EPM testing (Figure 3b). In contrast to the PL, loss of IL GIRK channel activity reduced immobility during the FST (Figure 3c). Conversely, there were no differences in correct responses and break points (Figure 3d) during progressive ratio tests. In the ASST, cre+ and cre- mice did not differ in their acquisition of lever training (Figure 3e), number of trials (Figure 3f-h), errors (Supplemental 4a-c), or omissions (Supplemental Table 4) during the visual cue, ED, or reversal test. These findings indicate that unlike the PL, IL GIRK-dependent signaling in male mice has little influence on affect and cognitive behaviors.

**Figure 3.**
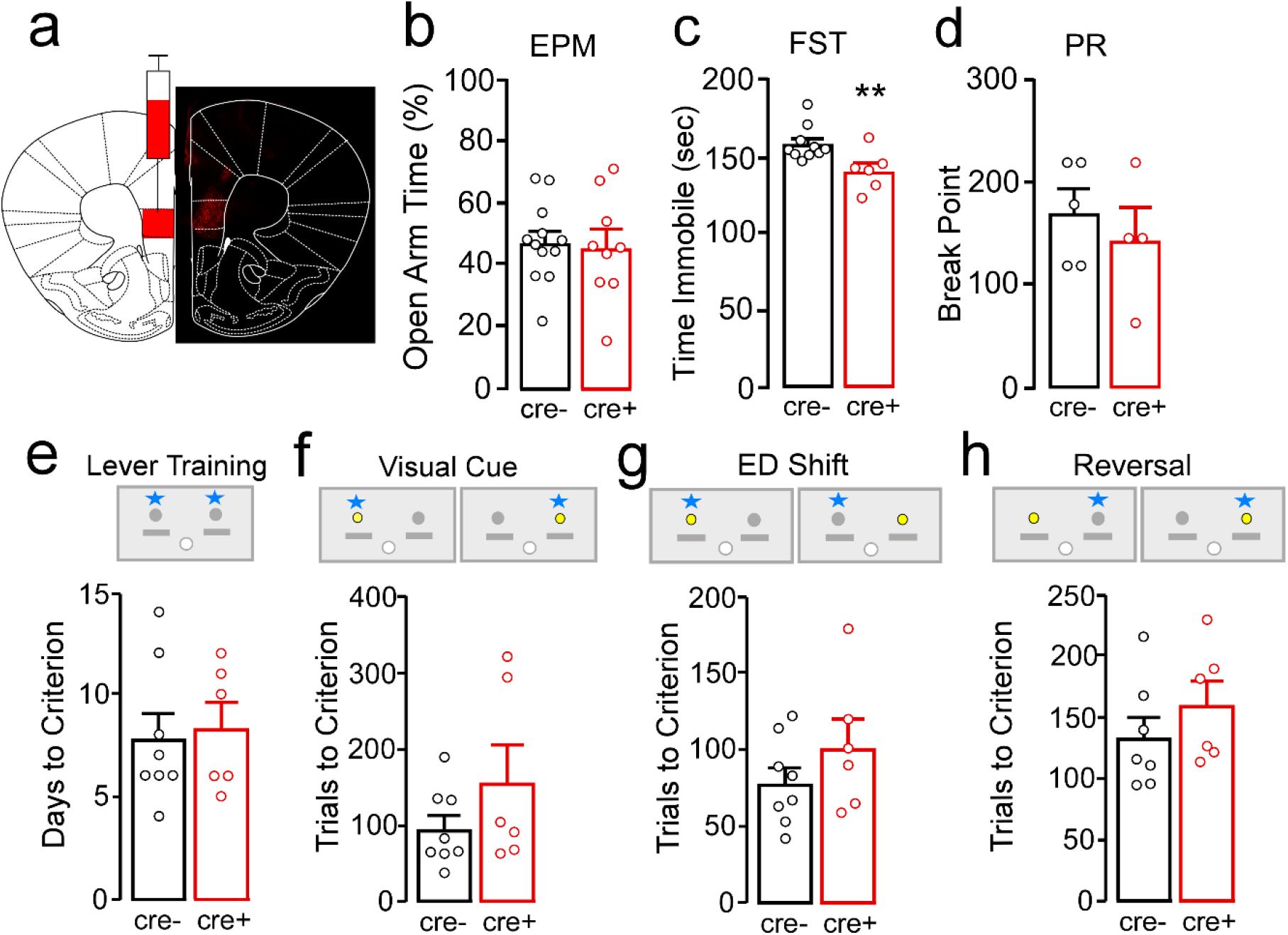
**a.** Schematic and representative image of AAV8-Cre-CAMKII viral targeting of the IL in Girk1^flox/flox^ male mice. **b.** Percent time spent in the open arm of the EPM did not differ in cre+ and cre- mice (t_(19)_= 0.25, p=0.80). **c.** Total time spent immobile during the FST was reduced in cre+ male mice compared to cre- (t_(14)_= 2.88, p=0.01). **d.** Break point (U=8.00, p=0.73) and correct responses (t_(7)_=0.38, p=0.72; data not shown) during progressive ratio testing did not differ between cre+ and cre- mice. **e-h.** Cre+ and cre- male mice took similar number of days to reach lever training criterion (**e**, t_(12)_= −1.21, p=0.25). Mice also took similar trials to reach criterion during the visual cue (**f**, t_(12)_= −1.31, p=0.22), ED shift (**g**, t_(12)_= −1.21, p=0.25), and the reversal test (**h**, t_(11)_= −1.04, p=0.32).

### Role of PL GIRK-dependent Signaling in Females

Intrinsic sex differences in mPFC physiology and function may have translational significance regarding resilience or susceptibility to pathological disorders (72–74). Past studies have identified sex-dependent differences in GIRK-dependent signaling (42), therefore we next examined whether loss of GIRK1 similarly impacts female behavior. Whole-cell recordings in Girk1^flox/flox^ female mice showed that similar to PL pyramidal neuron GIRK KO male mice (41, 75), I_Baclofen_ is reduced in cre+ versus cre- pyramidal neurons (Figure 4a). Unlike males, however, cre+ and cre- females had no difference in open arm time during the EPM (Figure 4b), time spent immobile in the FST (Figure 4c), or number of correct responses or break point (Figure 4d) during a progressive ratio. Moreover, no differences were observed between cre+ and cre- females across any measure related to the ASST, including days to reach lever training criteria (Figure 4e), or number of trials (Figure 4f-h), errors (Supplemental 5a-c), or omissions (Supplemental Table 5) to reach criterion during the visual cue, ED Shift, or reversal test. These data indicate that although GABA_B_-GIRK signaling in PL pyramidal neurons is present in males and females, this signaling mechanism plays a distinctly different role in regulation of PL function.

**Figure 4.**
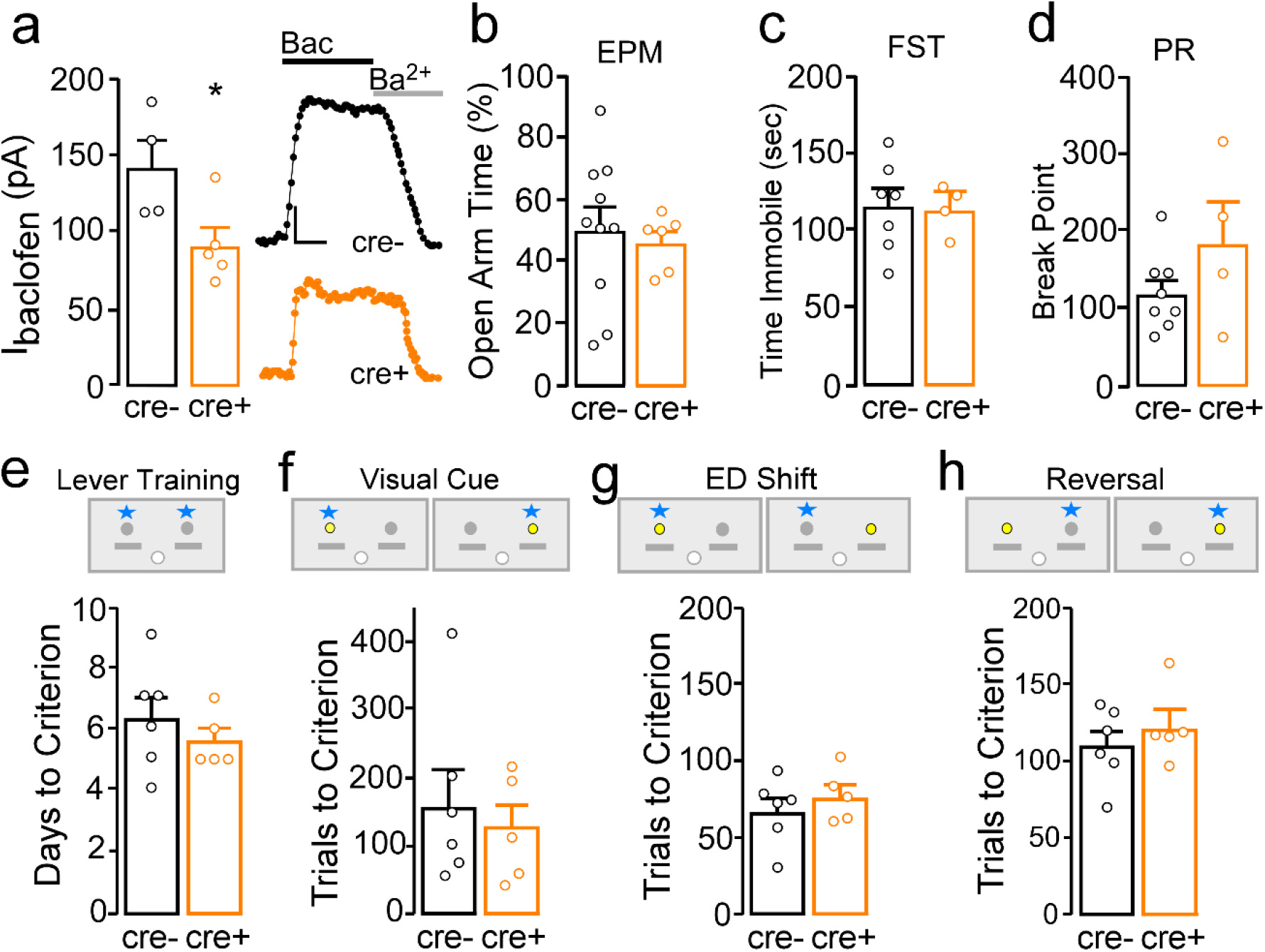
**a.** I_baclofen_ in Girk1^flox/flox^ female mice was significantly reduced in cre+ (n=5/N=4) L5/6 PL pyramidal neurons compared to cre- controls (t_(7)_=2.48, p=0.04; n=4/N=3). **b.** Percent open arm time in the EPM was similar in cre+ and cre- female mice (t_(14)_=0.38, p=0.71). **c.** Total time spent immobile during the forced swim test was similar between cre+ and cre- female mice (t_(8)_=0.14, p=0.89). **d.** Break point (U=2.50, p=0.07) nor correct responses (t_(8)_=-1.35, p=0.22; data not shown) did not differ in cre+ and cre- females. **e-h.** Cre+ and cre- female mice took similar number of days to reach lever training criterion (**e**, t_(9)_=0.84, p=0.42), trials to reach criterion during the visual cue test (**f**, t_(9)_=0.49, p=0.64), the ED Shift (**g**, t_(9)_=-0.79, p=0.45), and the reversal test (**h**, t_(9)_=-0.81, p=0.44).

### Role of PL GIRK-dependent signaling in chronic stress-induced deficits in cognitive flexibility

Chronic stress-induced mPFC dysfunction is thought to reflect imbalances in excitation:inhibition, however the mechanism through which this occurs has yet to be identified (76). Using a previously established mouse model of CUS (59), we determined if CUS promotes enduring changes in PL pyramidal GABA_B_-GIRK signaling (Figure 5a). Male mice exposed to 4 weeks of CUS showed a significant reduction in I_Baclofen_ (Figure 5b) and I_ML-297_ (Figure 5c) compared to naïve controls. In contrast, exposure to a similar regimen of CUS in females did not alter I_Baclofen_ (Figure 5d).

**Figure 5.**
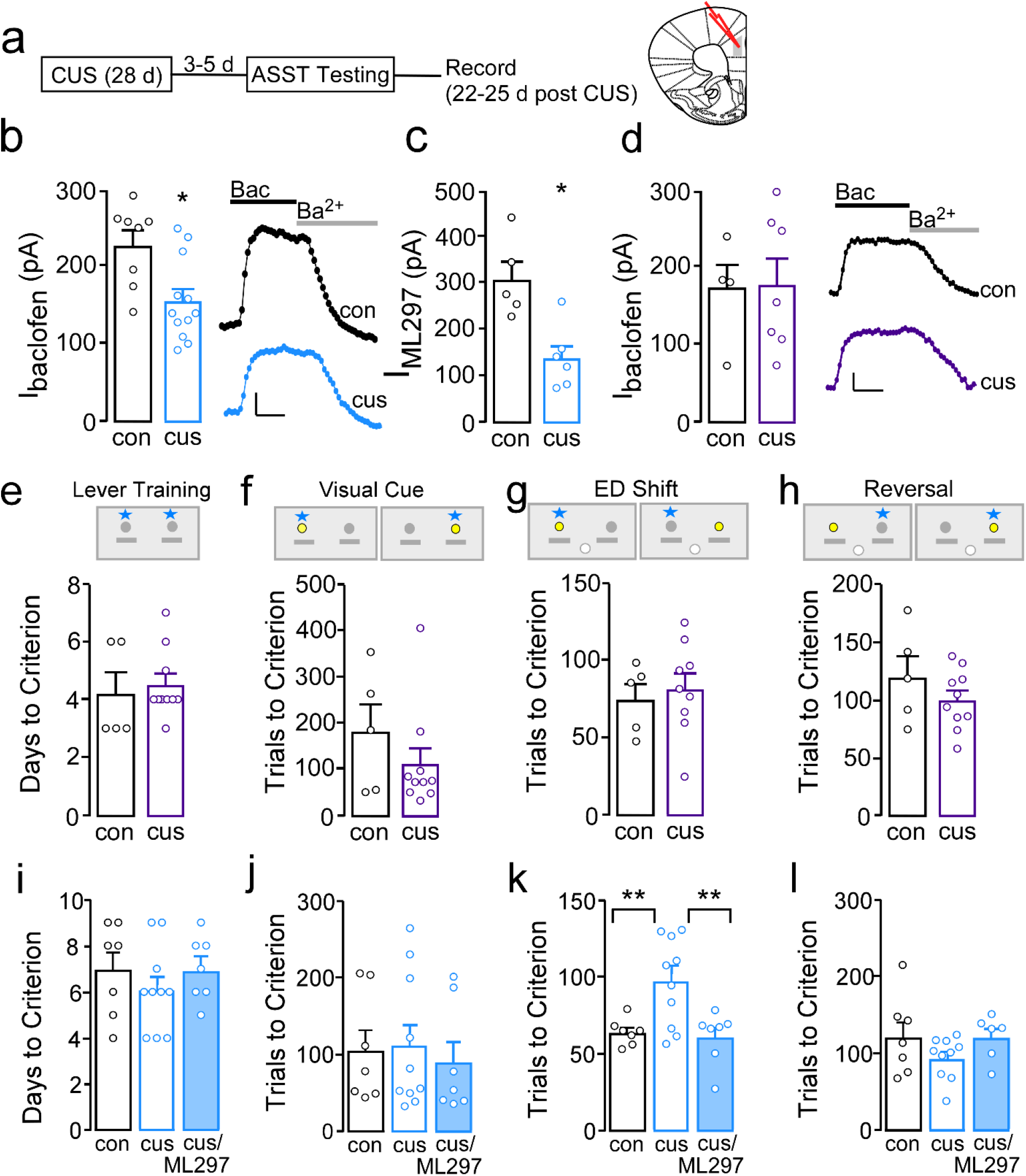
**a.** Timeline of CUS, behavioral testing, and electrophysiology recordings. **b.** I_baclofen_ in PL L5/6 was reduced in CUS exposed male mice compared to CUS naïve (con) (t_(18)_=3.04, p=0.007; con n=8/N=6; CUS n=12/N=9). **c.** I_ML297_ in PL L5/6 pyramidal neurons was significantly reduced in CUS male mice compared to con (t_(9)_=3.67, p=0.005; con n=5/N=3; CUS n=6/N=4). **d.** I_baclofen_ did not differ in PrL L5/6 PYR from CUS versus Con female mice (t_(9)_=-0.17, p=0.87; con n=4/N=4; CUS n=7/N=6). **e.** Female mice previously exposed to four weeks of CUS required a similar number of days to reach lever training criterion (t_(13)_= −0.41, p=0.69). **f-h.** There was also no difference in trials to reach criterion between the two groups during the visual cue test (**f**, t_(13)_=1.08, p=0.30), the ED shift (**g**, t_(12)_=-0.41, p=0.69), or during the reversal test (**h**, t_(13)_=1.16, p=0.27). **i.** There were no differences between the three groups of male mice in days to reach lever training criteria (F_(2, 21)_=0.74, p=0.49). **j.** No differences were observed in trials to reach criterion during the visual cue test (F_(2, 21)_=0.16, p=0.85). **k.** During the ED Shift, CUS male mice required more trials than control mice (p<0.01) and compared to CUS that received ML297 (p<0.01) whereas control mice and CUS mice that received ML297 did not differ (F_(2,21)_= 7.40, p=0.004). **l.** No differences was observed in trials to reach criteria for the reversal test (F_(2,20)_=1.64, p=0.22). *p<0.05, **p<0.01. Scale bars: 50pA x 100sec.

We previously demonstrated that our CUS model recapitulates a range of psychiatric-related phenotypes including increased anhedonia- and anxiety-like behavior (59), and reduced coping strategies in male mice. However, the impact of this regimen on cognitive function in C57BL/6J mice (a strain notoriously resilient to stress-induced cognitive deficits) (77), as well as in females was untested. Here we find that unlike males, female mice exposed to CUS do not exhibit changes in EPM open arm time or break points, however do show reduced time immobile during the FST time (Supplemental Figure 6a-c). Moreover, aligning with a lack of CUS exposure on baclofen-induced PL currents, CUS exposure in females did not impact performance on any ASST measures including days to lever train (Figure 5e) or on trials (Figure 5f-h), errors (Supplemental Figure 6d-f), or omissions (Supplemental Table 6) to reach criteria during each of the three tests.

In contrast, initial cohorts of *male mice* indicated that CUS exposure impaired performance in the ED Shift. Thus, in subsequent groups, male mice were divided into three conditions: stress naïve (con), CUS, or CUS injected with ML297 prior to the ED Shift (CUS/ML297). No differences were observed across condition for days to reach lever training criteria (Figure 5i) or trials (Figure 5j), errors (Supplemental Figure 7a) or omissions (Supplemental Table 7) during the visual cue test. During the ED test, there was a significant difference in trials (Figure 5k), but not errors (Supplemental 7b), to reach criteria, with post-hoc comparisons showing CUS mice required more trials than con (p=0.006) and CUS/ML297 (p=0.008), while con and CUS/ML297 did not differ (p=0.78). A significant difference in omissions was detected; CUS/ML297 mice had higher omissions compared to con (p=0.005) and CUS mice (p=0.001; Supplemental Table 7). During reversal testing, CUS/ML297, CUS, and con mice showed no distinction in trials (Figure 5l), errors (Supplemental 7c), or omissions (Supplemental Table 7) to criteria. Together, these findings indicate that CUS differentially impacts cognitive flexibility and PL physiology in males versus females and that deficits in male mice are driven in part by reductions in GABAB-GIRK signaling that can be rescued by systemic activation of these channels.

## DISCUSSION

### Impact of GIRK1 Ablation on Affect and Cognition in Males

The neurophysiological substrates of psychiatric and stress-related disorders are poorly understood. However, an emerging theory is that imbalances in PFC cellular excitation:inhibition can give rise to abnormalities in both affect and cognition, and thus represents a unifying substrate for shared pathology across neuropsychiatric disorders. Our past work has shown that excitation:inhibition of PL pyramidal neurons is highly regulated by GIRK channels, however, the contribution of PL GIRK channels on regulation of affect and cognition, and whether disruption in PL GIRK signaling contributes to stress-related maladaptive behavior has never been examined. Constitutive knockout of GIRK1 or GIRK2 reduces anxiety-like behavior (52, 53), whereas global reductions in GIRK channel activity promote depression-resistant phenotypes (78) and increase motivation for appetitive rewards (52). Here, we found that loss of PL pyramidal neuron GIRK channels increases EPM open arm time, increases FST time immobile, and may augment motivation, suggesting that PL GIRK1-dependenet signaling is a primary effector of behaviors observed in knockout mice, and that the resulting increase in PL pyramidal neuron excitation alters neuronal processing necessary for normal behavioral responses in these tests (e.g., less time in an EPM open arm) (79–83).

Cognitive processes associated with working memory are reliant, although not exclusively, on coordinated PL activity (84–87). Here we find that GIRK PL ablation produces a consistent reduction in working memory using a forced alternation model. Assessment of more complex processes using the ASST showed an impaired performance in cre+ during the visual cue test in males. PL lesion was previously shown to disrupt performance in the ED Shift but not visual cue test (14, 34), thus our observations were unexpected. Given that loss of GIRK channel activity did not alter the ability to acquire a visual cue-based learning task, it is unlikely that this deficit reflects general attention deficits to a cue or reduced visual acuity (88). Rather, loss of GIRK1 alters PL function in a manner where the addition of a visual cue contingency not present during lever training is sufficient to act as an attentional shift. In support, experiments involving response-to-cue shifts showed that cre+ mice also performed worse when the contingency was changed from lever training to response (attend to one lever only), and that these impairments persisted during a subsequent response-to-cue ED Shift. Thus, it is likely that increased/disorganized firing following loss of PL pyramidal GIRK underlies deficits in working memory and cognitive flexibility.

The PL and IL have differences in both connectivity and contributions to behavior (24, 69, 70, 89). Importantly, GABA_B_-GIRK signaling has been demonstrated in IL L5/6 pyramidal neurons of male mice (41, 42) however the current experiments suggest that this signaling is only critical for FST time immobile. These effects on affect-related behaviors is not unfounded, as PFC dysfunction can have a divalent impact on affective behavior in humans and rodents depending on the downstream target and/or pyramidal sub-type (14, 24, 69, 70, 90–93).

### Impact of CUS on GABA_B_-GIRK and Cognitive Flexibility

Similar to cognitive inflexibility in humans (9, 94, 95), exposure to CUS impairs performance in the ED Shift test in male mice. Deficits in flexibility aligned with a reduction in GABA_B_-GIRK signaling, highlighting a role in CUS-induced PL dysfunction. However, unlike GIRK1 knockouts, deficits in flexibility following CUS were specific to the ED Shift as has previously been shown following restraint stress in male rats (96). Similar to PL lesion studies (14, 34), CUS exposure may impair ED Shift responding rather than visual cue responding as it likely has more global effects than just reducing PL pyramidal neuron GIRK-dependent signaling. As converging lines of evidence show that exposure to unpredictable stressors more negatively impacts cognitive control in humans and rodents (9, 94, 95, 97, 98), the ability of systemic ML297 to rescue this deficit has significant impact on future therapeutics aimed at treating cognition-related deficits produced by stress and other psychiatric conditions.

### PL PYR GIRK1 Knockout and CUS Exposure in Females

Sex differences in susceptibility to anxiety, mood, and other neuropsychiatric disorders has been established in humans (99–102). Therefore, the influence that mPFC physiology and function has on behavior in both males and females has significant translational value for identifying both susceptibility and treatment options. While we replicated findings showing that GABA_B_-GIRK signaling is present in female PL pyramidal neurons (42), ablation did not impact affect or cognitive flexibility, suggesting that this signaling is not as critical for PL function in females. Although not examined, it is unlikely estrous stage played a role in the lack of effect, as GABA_B_-GIRK signaling and rheobase does not differ across stage (42). We also found that our regimen of CUS was more impactful in males, as females exhibited no changes in EPM, progressive ratio, cognitive flexibility, or GABA_B_-GIRK signaling. This may reflect resiliency to stress, as prior studies have shown chronic restraint stress impairs performance in males but either does not influence or enhances performance in females in a variety of tasks, including behavioral flexibility (103–111). Females also have attenuated mPFC dendritic and plasticity changes following chronic stress compared to males (112–114). Conversely, the lack of CUS effects in females likely reflects the type of stress exposure. Rather than unpredictable stress models (e.g. restraint, forced swim) which have been optimized for male rodents, social stressors have increased ethological validity for female rodents (115) and social stress has been shown to impair reversal learning during a set-shift task in females (116) and alter mPFC plasticity in female rats (117).

Collectively, we highlight the importance of GIRK-dependent signaling in male PL pyramidal neurons in the regulation of both affect and cognition and demonstrate that PL GIRK channels in males are targets of stress. Further, our data, along with past studies (51, 53), suggests that drugs that target GIRK channels and related G protein inhibitory signaling may serve as clinically relevant therapeutic strategies to treat both cognitive and affect-related symptoms.

## Acknowledgements

These studies were supported by funding from the Brain and Behavior Research Foundation (#26299; MH), Marquette University Regular Research Grant (MH), the Charles E Kubly Mental Health Research Foundation at Marquette University (MH), and NIH grants DA034696 and AA027544 (KW).

## Disclosure

The authors have no biomedical financial interests or potential conflicts of interest.

---

**Supplemental Figure 1.**
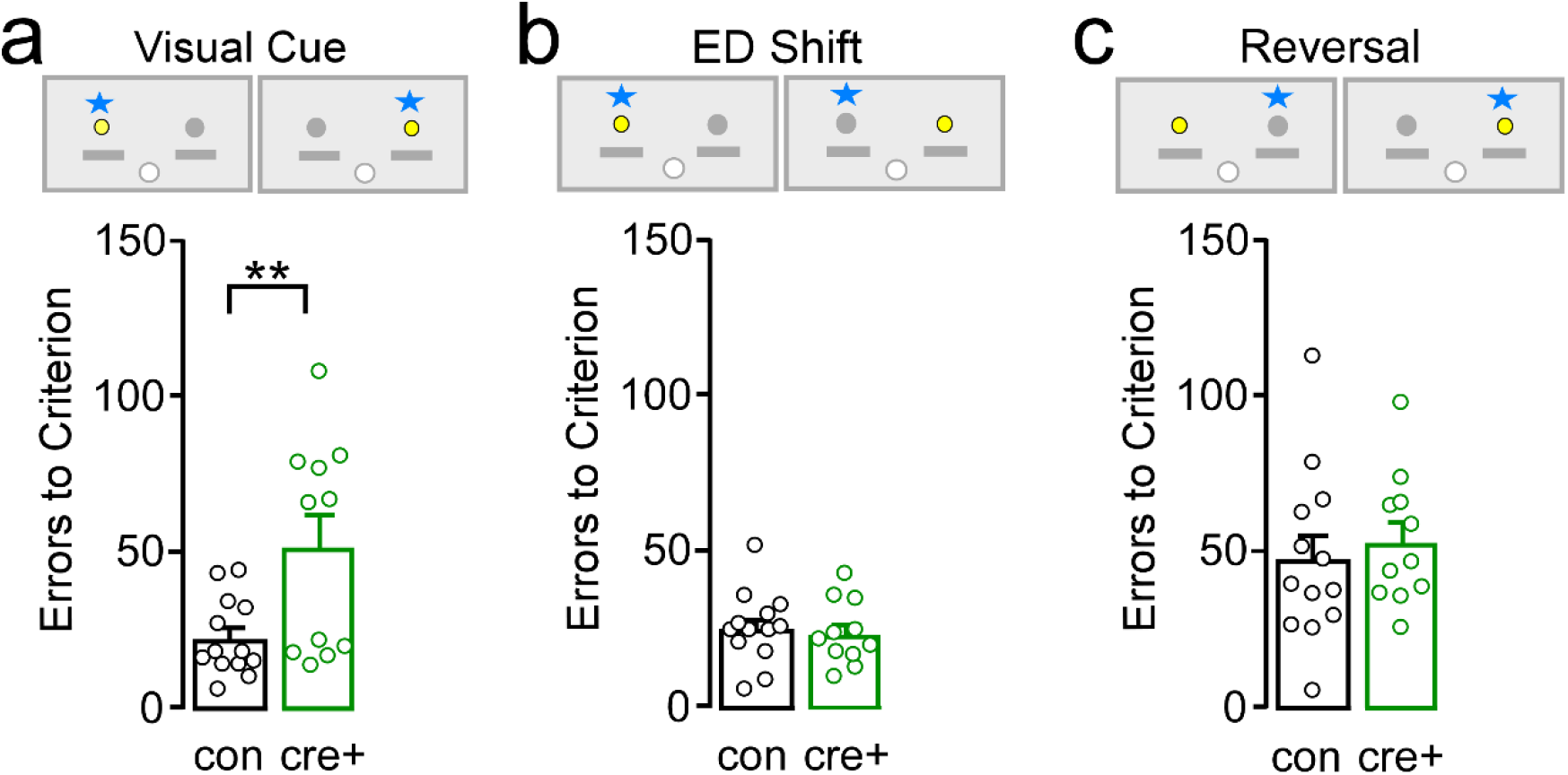
**a.** Numbers of errors prior to reaching criterion of ten correct responses in a row during the visual cue test. Cre+ had more errors compared to cre- mice (t_(22)_=-2.91, p=0.008). **b.** Number of errors taken to reach criterion during the ED Shift test. There were no differences between cre- controls and cre+ male mice (t_(22)_= 0.38, p=0.71). **c.** Errors to reach criterion during the reversal test. There were no differences between control male mice or cre+ male mice (t_(22)_= −0.55, p=0.59). ** p<0.01.

**Supplemental Figure 2.**
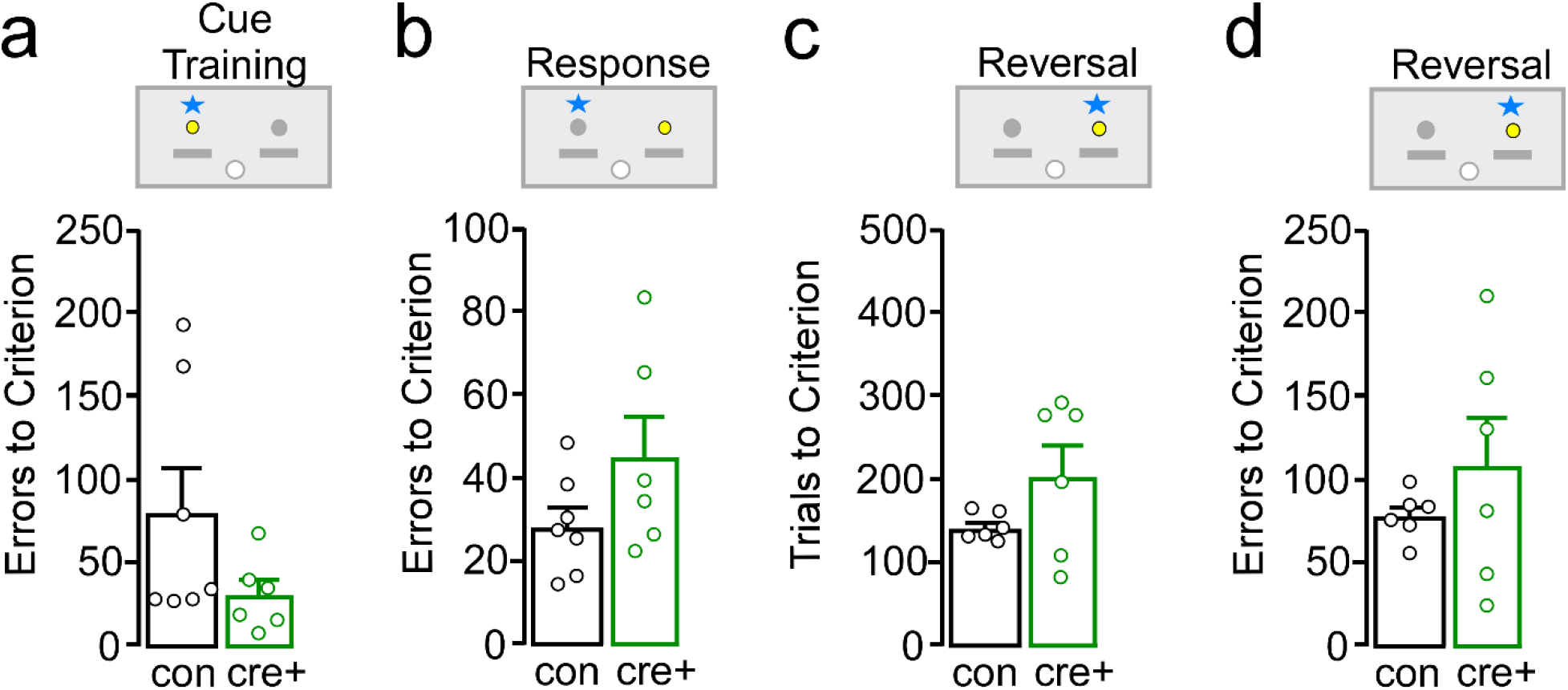
**a.** Numbers of errors prior to reaching criterion of ten correct responses in a row during the visual cue training. Cre- controls had had similar number of errors compared to cre+ mice (t_(11)_=1.59, p=0.14). **b.** Number of errors taken to reach criterion during the ED Shift test. There were no differences between controls (cre-) and cre+ male mice (t_(11)_=-1.61, p=0.14). **c.** There were no differences in trials to reach criterion during the reversal test between cre- and cre+ mice (t_(10)_=-1.65, p=0.13). **d.** Errors to reach criterion during the reversal test. There were no differences between control male mice and cre+ male mice (t_(10)_=-1.02, p=0.33).

**Supplemental Figure 4.**
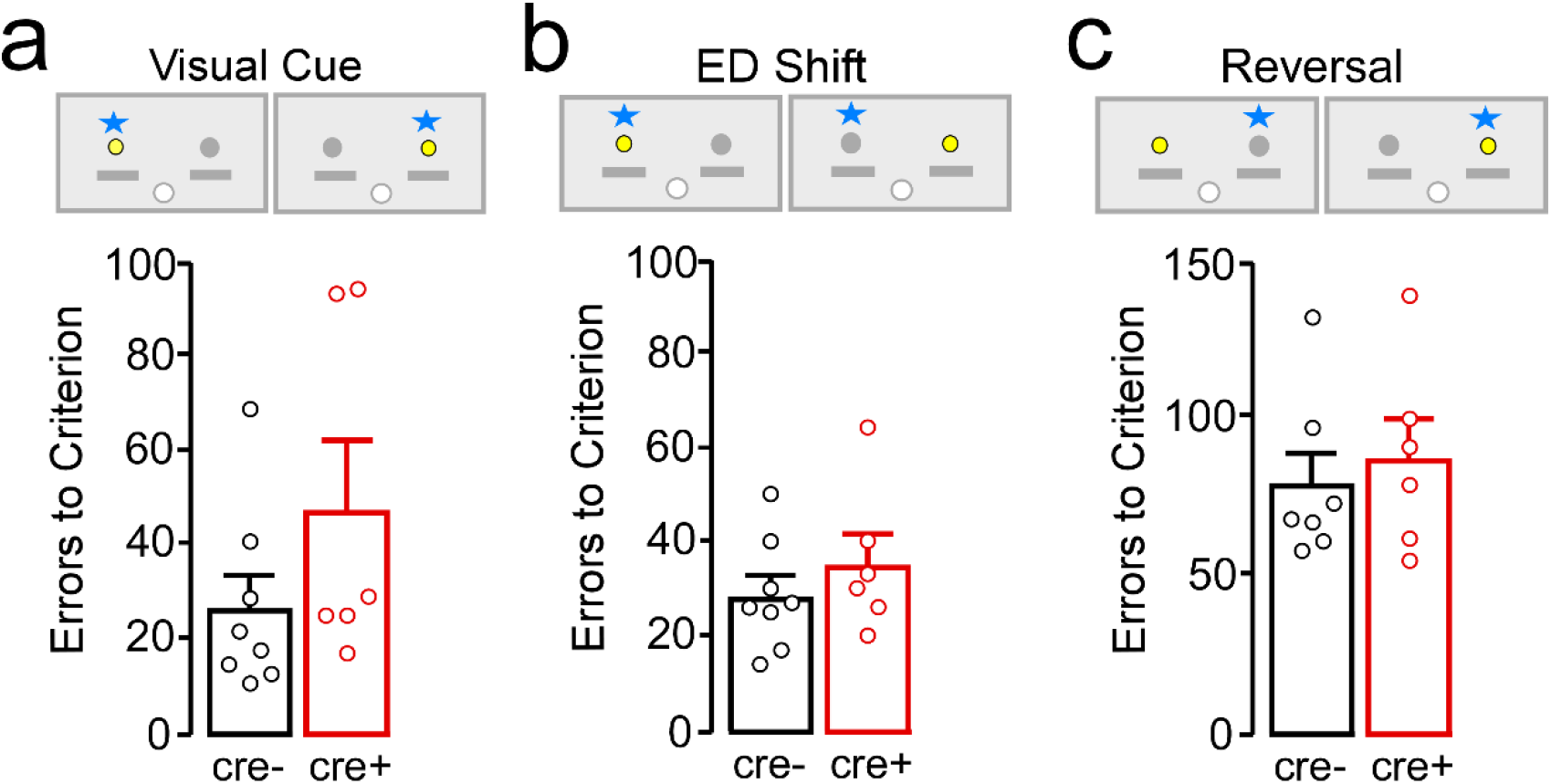
**a.** Number of errors made prior to reaching criterion during the visual cue test in control male mice (cre-) compared to male mice with bilateral infusion of AAV-cre into the IL. There were no differences between the two groups (t_(12)_= −1.40, p=0.19). **b.** There was no difference in number of errors made prior to reaching criterion during the ED Shift test between control cre- mice and IL cre+ mice (t_(12)_=-0.95, p=0.36). **c.** The number of errors made during the reversal test was similar for cre- mice compared to IL cre+ male mice (t_(11)_=-0.52, p=0.61).

**Supplemental Figure 3.**
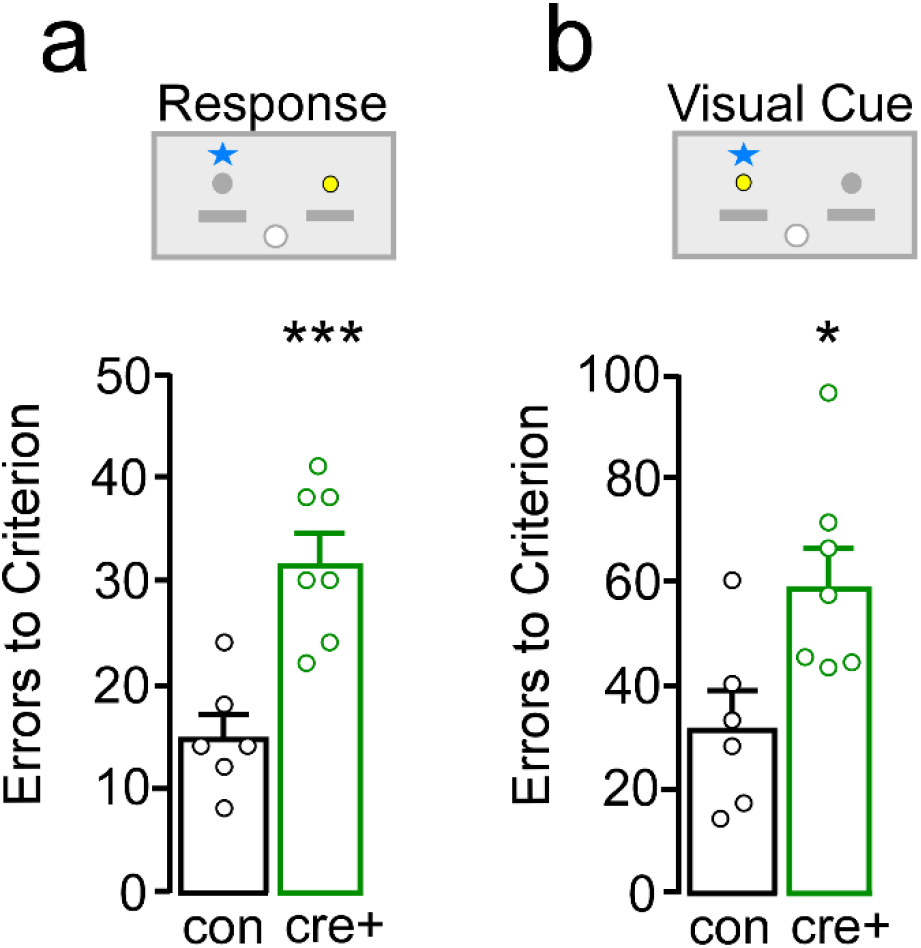
**a.** Number of errors prior to reaching criterion of ten correct responses in a row during the response test. Cre+ mice had significantly more errors to criterion compared to cre- controls (t_(11)_= −4.61, p<0.001). **b.** Number of errors taken to reach criterion during the visual cue test. Cre+ male mice had significantly more errors to criterion compared to cre- controls (t_(11)_= −2.69, p=0.02). * p<0.05; *** p<0.001.

**Supplemental Figure 5.**
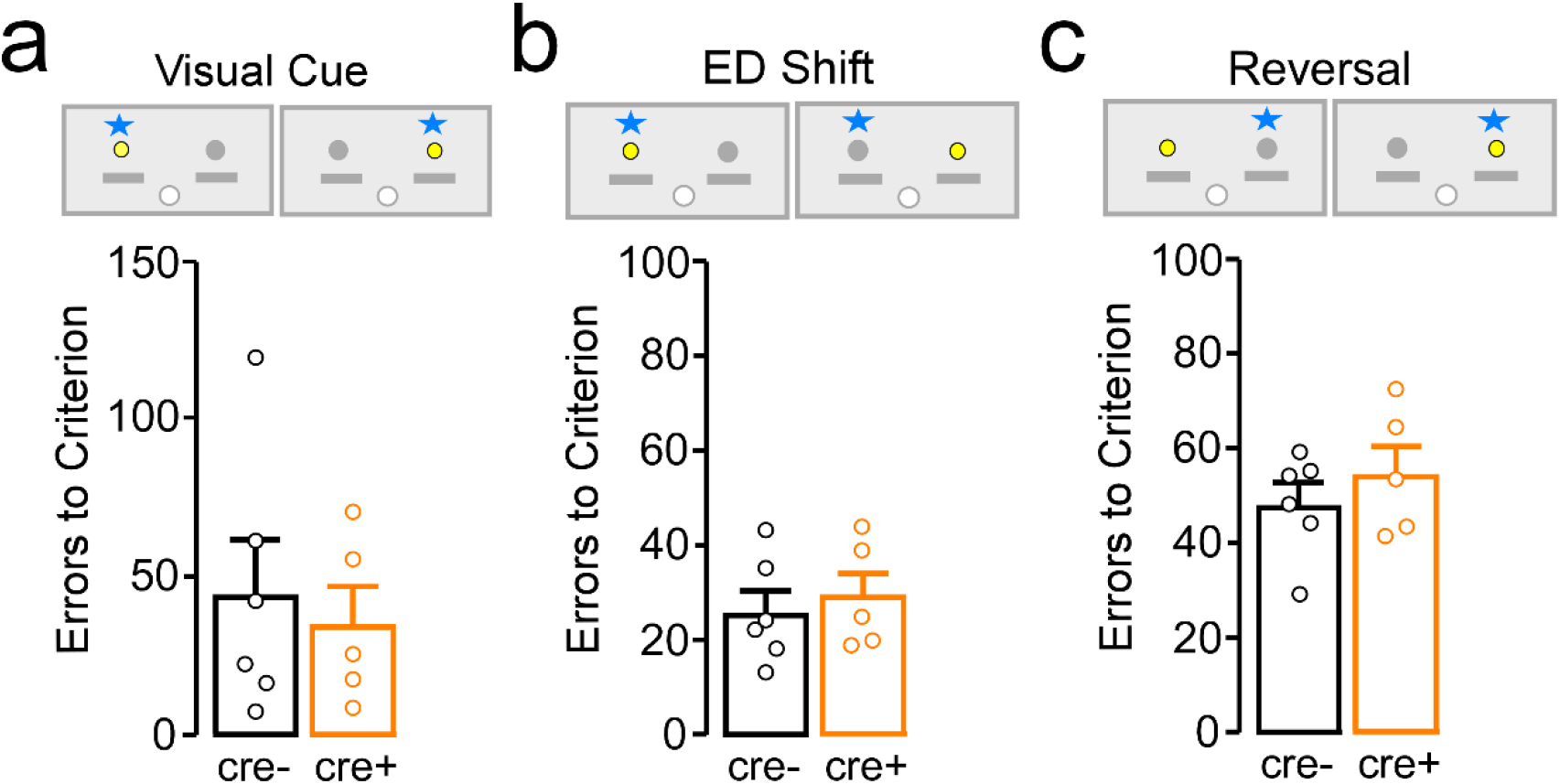
**a.** Errors to reach criterion during the visual cue test in control female mice (cre-) compared to female mice with bilateral infusion of AAV-cre into the PL were similar between the two groups (t_(9)_=0.44, p=0.67). **b.** There were no differences between female cre- and cre+ mice in number of errors to reach criterion during the extradimensional shift test (t_(9)_=- 0.52, p=0.61). **c.** The number of errors made during the reversal test was similar for control cre- female mice compare to cre+ female mice (t_(9)_=-0.89, p=0.40).

**Supplemental Figure 6.**
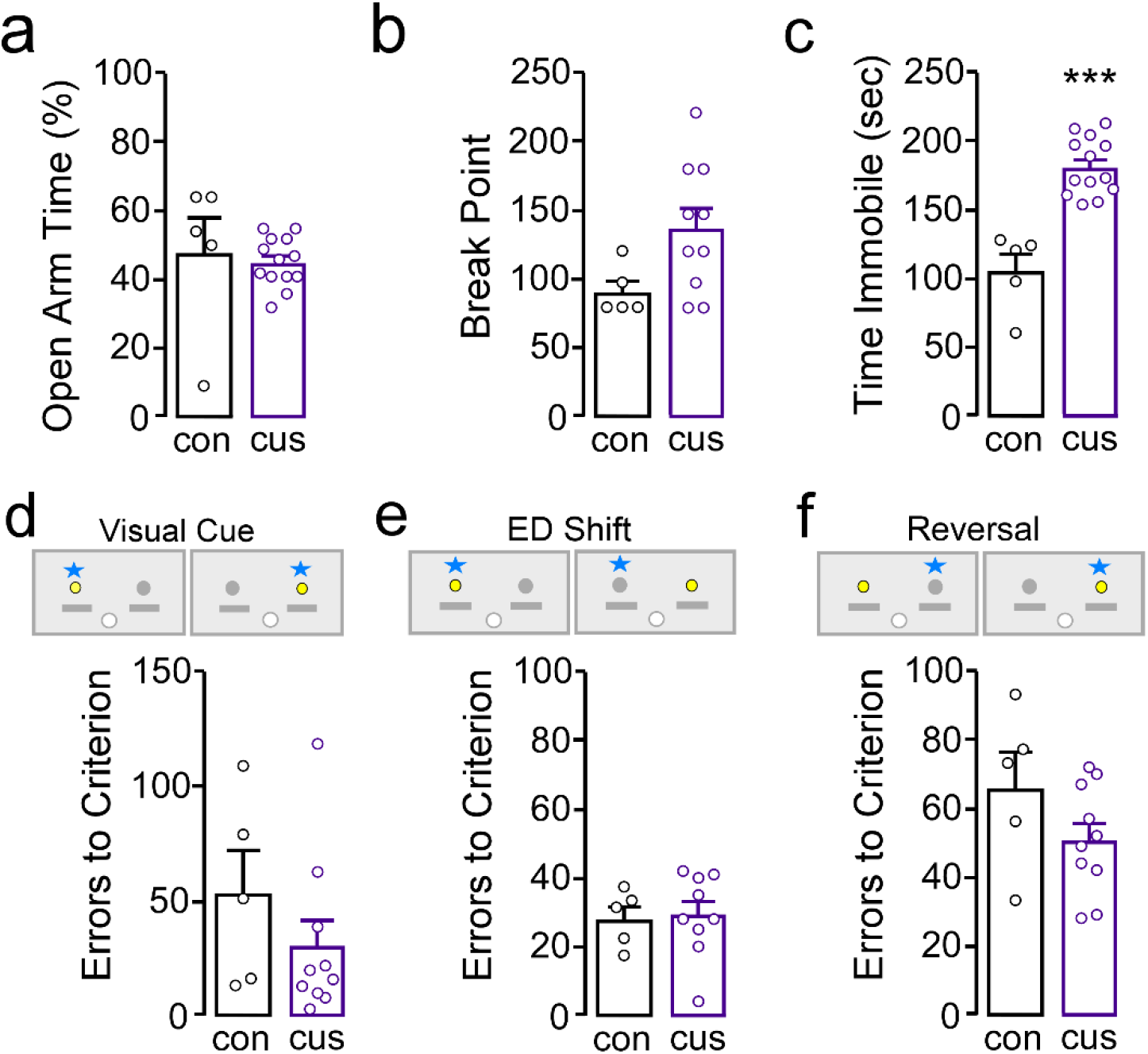
**a.** Percent time in the open arm during the elevated plus maze. Control female mice spent similar percent time in the open arm compared to female mice previously exposed to four week of CUS (t_(16)_=0.42, p=0.68). **b.** During a progressive ratio test for an ensure reward, there was a trend towards increased break points in female mice exposed to CUS compared to controls (U=9.50, p=0.06). **c.** Female mice previously exposed to CUS spent significantly more time immobile in the FST compared to controls (t_(16)_=-6.21, p<0.001). **d.** Female mice exposed previously to four weeks of CUS had similar number of errors to reach criterion during a visual cue test (t_(13)_=1.14, p=0.28). **e.** During the extradimensional shift test, female control mice had similar errors to criterion compared to CUS mice (t_(12)_=-0.20, p=0.85). **f.** During the reversal test, female control mice had similar errors to reach criterion compared to CUS mice (t_(13)_=1.54, p=0.15). ***p<0.001

**Supplemental Figure 7.**
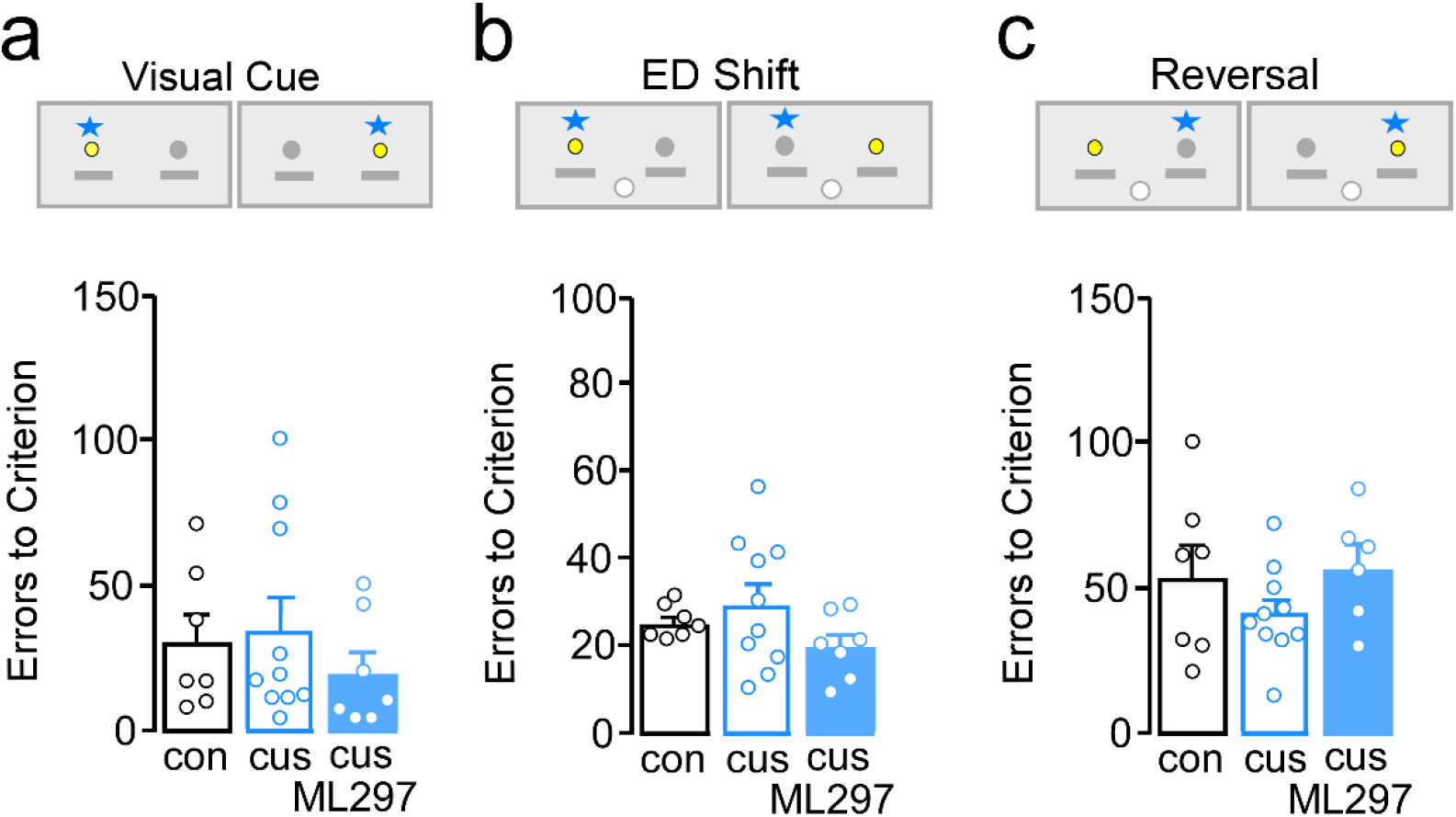
**a.** The three groups of male mice (con, CUS, CUS that would receive ML297 prior to ED test) had similar number of errors to reach criterion during the visual cue test (F_(2, 21)_=0.61, p=0.55). **b.** The three groups did not differ on errors to reach criterion during the extradimensional shift test (F_(2,21)_= 1.62, p=0.22). **c.** During the reversal test, the three groups had similar number of errors to reach criterion (F_(2,20)_=1.31, p=0.29).

**Supplemental Table 1.** Number of omissions for cre- control male mice and cre+ male mice during visual cue test, ED Shift, and reversal test. There were no differences in number of omissions during the visual cue (U=46.50, p=0.13), ED shift (U=71.00, p=1.00), and reversal test (t_(22)_=-1.36, p=0.19). Numbers denote mean ± standard error of mean.

|  | Visual Cue | ED Shift | Reversal |
| --- | --- | --- | --- |
| **Control (cre-)** | 1.31 ± 0.75 | 1.54 ± 0.46 | 2.08 ± 0.71 |
| **Cre+** | 4.18 ± 1.59 | 3.00 ± 1.63 | 11.09 ± 7.21 |

**Supplemental Table 2.**
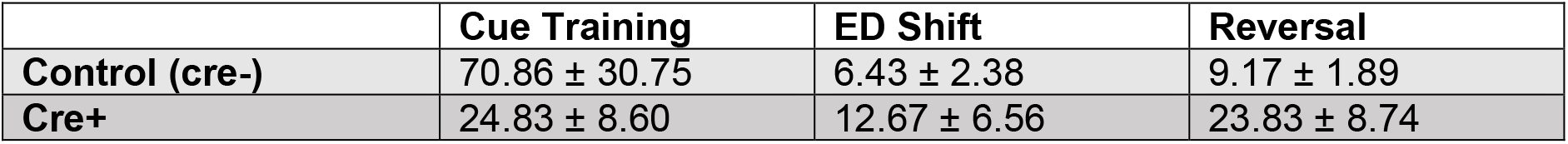
Number of omissions for cre- control male mice and cre+ male mice for visual cue training, ED Shift, and reversal test. There were no differences in number of omissions during the visual cue training (t_(11)_=1.34, p=0.21), ED Shift (t_(11)_=-0.95, p=0.36), or reversal test (t_(10)_=-1.64, p=0.13) for control (cre-) and AAV (cre+) male mice. Numbers denote mean ± standard error of mean.

**Supplemental Table 3.**
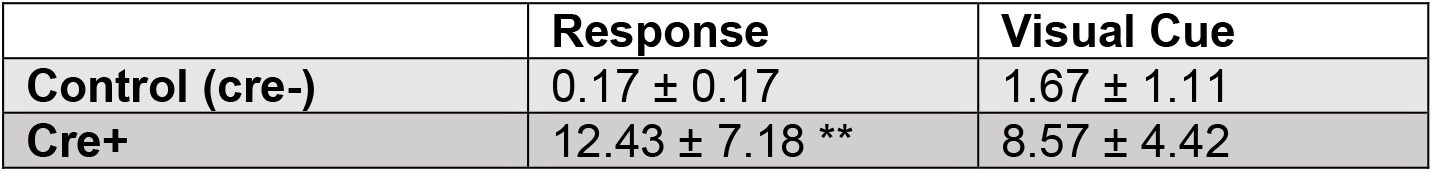
Number of omissions for cre- controls and cre+ during a response test and a visual cue test. Cre+ mice had significantly more omissions compared to cre- controls during the response test (U=3.50, p=0.008) however there were no differences in number of omissions during the visual cue test (t_(11)_= −1.40, p=0.19). Numbers denote mean ± standard error of mean. ** p<0.01.

**Supplemental Table 4.** Number of omissions for cre- control male mice and IL cre+ male mice during visual cue test (U=20.00, p=0.66), ED shift test, and reversal test. There were no differences in number of omissions during the visual cue, ED Shift (t_(12)_=-0.93, p=0.37), and reversal test (t_(11)_=0.05, p=0.95). Numbers denote mean ± standard error of mean.

|  | Visual Cue | ED Shift | Reversal |
| --- | --- | --- | --- |
| **Control (cre-)** | 0.50 ± 0.19 | 1.00 ± 0.38 | 4.29 ± 1.57 |
| **Cre+** | 2.17 ± 1.60 | 2.50 ± 1.82 | 4.17 ± 1.08 |

**Supplemental Table 5.** Number of omissions for cre- control female mice and cre+ female mice during visual cue test, ED shift test, and reversal test. There were no differences in number of omissions during the visual cue (U=13.00, p=0.79), ED Shift (t_(9)_=-0.63, p=0.55), and reversal test (t_(9)_=0.11, p=0.91). Numbers denote mean ± standard error of mean.

|  | Visual Cue | ED Shift | Reversal |
| --- | --- | --- | --- |
| **Control (cre-)** | 4.33 ± 2.40 | 1.50 ± 0.56 | 8.33 ± 4.91 |
| **Cre+** | 3.80 ± 2.40 | 2.00 ± 0.55 | 7.60 ± 3.82 |

**Supplemental Table 6.** Number of omissions during visual cue, ED shift, and reversal test of the ASST for control female mice compared to female mice previously exposed to 4 weeks of chronic unpredictable stress (CUS). There were no differences between the two groups for the visual cue test (t_(13)_=2.13, p=0.053), the ED shift test (U=22.50, p=0.93), or the reversal test (U=11.00, p=0.09).

|  | Visual Cue | ED Shift | Reversal |
| --- | --- | --- | --- |
| **Control** | 6.00 ± 3.51 | 0.40 ± 0.40 | 4.60 ± 1.57 |
| **CUS** | 0.80 ± 0.39 | 0.33 ± 0.24 | 1.40 ± 0.40 |

**Supplemental Table 7.** Number of omissions during visual cue, ED Shift, and reversal test of the ASST for control male mice compared to male mice previously exposed to 4 weeks of CUS and those exposed to CUS that also received ML297 on the ED Shift test day. There were no differences in omissions on visual cue test (H_(2)_=0.56, p=0.76) or reversal test (F_(2,20)_=0.61, p=0.55), however during the ED Shift there was a significant difference between the three groups (F_(2,21)_=8.57, p=0.002) as CUS mice that received ML297 had significantly more omissions than control mice (**p<0.01) and CUS mice without ML297 (++p<0.01).

|  | Visual Cue | ED Shift | Reversal |
| --- | --- | --- | --- |
| **Control** | 1.71 ± 1.29 | 1.00 ± 0.85 | 3.71 ± 2.03 |
| **CUS** | 1.50 ± 0.79 | 1.40 ± 0.88 | 2.40 ± 0.64 |
| **CUS / ML297** | 0.29 ± 0.18 | 11.14 ± 3.44**++ | 1.67 ± 0.88 |

## Notes

### Competing Interest Statement

The authors have declared no competing interest.

## References

1. Bale TL. Sensitivity to stress: dysregulation of CRF pathways and disease development. Horm Behav. 2005;48(1):1–10.

2. Kendler KS, Karkowski LM, Prescott CA. Stressful life events and major depression: risk period, long-term contextual threat, and diagnostic specificity. J Nerv Ment Dis. 1998;186(11):661–9.

3. Kendler KS, Karkowski LM, Prescott CA. Causal relationship between stressful life events and the onset of major depression. Am J Psychiatry. 1999;156(6):837–41.

4. Moghaddam B, Javitt D. From revolution to evolution: the glutamate hypothesis of schizophrenia and its implication for treatment. Neuropsychopharmacology. 2012;37(1):4–15.

5. Marazziti D, Consoli G, Picchetti M, Carlini M, Faravelli L. Cognitive impairment in major depression. Eur J Pharmacol. 2010;626(1):83–6.

6. Remijnse PL, van den Heuvel OA, Nielen MM, Vriend C, Hendriks GJ, Hoogendijk WJ, et al. Cognitive inflexibility in obsessive-compulsive disorder and major depression is associated with distinct neural correlates. PLoS One. 2013;8(4):e59600.

7. Scult MA, Knodt AR, Swartz JR, Brigidi BD, Hariri AR. Thinking and Feeling: Individual Differences in Habitual Emotion Regulation and Stress-Related Mood are Associated with Prefrontal Executive Control. Clin Psychol Sci. 2017;5(1):150–7.

8. Dajani DR, Uddin LQ. Demystifying cognitive flexibility: Implications for clinical and developmental neuroscience. Trends Neurosci. 2015;38(9):571–8.

9. Schwabe L, Wolf OT. Stress prompts habit behavior in humans. J Neurosci. 2009;29(22):7191–8.

10. Zeng H, Lee TM, Waters JH, So KF, Sham PC, Schottenfeld RS, et al. Impulsivity, cognitive function, and their relationship in heroin-dependent individuals. J Clin Exp Neuropsychol. 2013;35(9):897–905.

11. Morilak DA, Frazer A. Antidepressants and brain monoaminergic systems: a dimensional approach to understanding their behavioural effects in depression and anxiety disorders. Int J Neuropsychopharmacol. 2004;7(2):193–218.

12. Negron-Oyarzo I, Aboitiz F, Fuentealba P. Impaired Functional Connectivity in the Prefrontal Cortex: A Mechanism for Chronic Stress-Induced Neuropsychiatric Disorders. Neural Plast. 2016;2016:7539065.

13. Gold JI, Treadwell M, Weissman L, Vichinsky E. An expanded Transactional Stress and Coping Model for siblings of children with sickle cell disease: family functioning and sibling coping, self-efficacy and perceived social support. Child Care Health Dev. 2008;34(4):491–502.

14. Holmes A, Wellman CL. Stress-induced prefrontal reorganization and executive dysfunction in rodents. Neurosci Biobehav Rev. 2009;33(6):773–83.

15. Strauss GP, Duke LA, Ross SA, Allen DN. Posttraumatic stress disorder and negative symptoms of schizophrenia. Schizophr Bull. 2011;37(3):603–10.

16. Strauss K, Vicari S, Valeri G, D’Elia L, Arima S, Fava L. Parent inclusion in Early Intensive Behavioral Intervention: the influence of parental stress, parent treatment fidelity and parent-mediated generalization of behavior targets on child outcomes. Res Dev Disabil. 2012;33(2):688–703.

17. Birrell JM, Brown VJ. Medial frontal cortex mediates perceptual attentional set shifting in the rat. J Neurosci. 2000;20(11):4320–4.

18. Bissonette GB, Martins GJ, Franz TM, Harper ES, Schoenbaum G, Powell EM. Double dissociation of the effects of medial and orbital prefrontal cortical lesions on attentional and affective shifts in mice. J Neurosci. 2008;28(44):11124–30.

19. Boettiger CA, D’Esposito M. Frontal networks for learning and executing arbitrary stimulus-response associations. J Neurosci. 2005;25(10):2723–32.

20. Murrough JW, Iacoviello B, Neumeister A, Charney DS, Iosifescu DV. Cognitive dysfunction in depression: neurocircuitry and new therapeutic strategies. Neurobiol Learn Mem. 2011;96(4):553–63.

21. Sullivan RM. Hemispheric asymmetry in stress processing in rat prefrontal cortex and the role of mesocortical dopamine. Stress. 2004;7(2):131–43.

22. Goldstein RZ, Volkow ND. Dysfunction of the prefrontal cortex in addiction: neuroimaging findings and clinical implications. Nat Rev Neurosci. 2011;12(11):652–69.

23. McKlveen JM, Morano RL, Fitzgerald M, Zoubovsky S, Cassella SN, Scheimann JR, et al. Chronic Stress Increases Prefrontal Inhibition: A Mechanism for Stress-Induced Prefrontal Dysfunction. Biol Psychiatry. 2016;80(10):754–64.

24. Dalley JW, Cardinal RN, Robbins TW. Prefrontal executive and cognitive functions in rodents: neural and neurochemical substrates. Neurosci Biobehav Rev. 2004;28(7):771–84.

25. Riga D, Matos MR, Glas A, Smit AB, Spijker S, Van den Oever MC. Optogenetic dissection of medial prefrontal cortex circuitry. Front Syst Neurosci. 2014;8:230.

26. Floresco SB, Block AE, Tse MT. Inactivation of the medial prefrontal cortex of the rat impairs strategy set-shifting, but not reversal learning, using a novel, automated procedure. Behav Brain Res. 2008;190(1):85–96.

27. Bissonette GB, Roesch MR. Neural correlates of rules and conflict in medial prefrontal cortex during decision and feedback epochs. Frontiers in behavioral neuroscience. 2015;9:266.

28. Yizhar O, Fenno LE, Prigge M, Schneider F, Davidson TJ, O’Shea DJ, et al. Neocortical excitation/inhibition balance in information processing and social dysfunction. Nature. 2011;477(7363):171–8.

29. Isaacson JS, Scanziani M. How inhibition shapes cortical activity. Neuron. 2011;72(2):231–43.

30. Sohal VS, Zhang F, Yizhar O, Deisseroth K. Parvalbumin neurons and gamma rhythms enhance cortical circuit performance. Nature. 2009;459(7247):698–702.

31. Kim H, Ahrlund-Richter S, Wang X, Deisseroth K, Carlen M. Prefrontal Parvalbumin Neurons in Control of Attention. Cell. 2016;164(1-2):208–18.

32. Gonzalez-Burgos G, Lewis DA. NMDA receptor hypofunction, parvalbumin-positive neurons, and cortical gamma oscillations in schizophrenia. Schizophr Bull. 2012;38(5):950–7.

33. Covington HE, 3rd, Lobo MK, Maze I, Vialou V, Hyman JM, Zaman S, et al. Antidepressant effect of optogenetic stimulation of the medial prefrontal cortex. J Neurosci. 2010;30(48):16082–90.

34. Enomoto T, Tse MT, Floresco SB. Reducing prefrontal gamma-aminobutyric acid activity induces cognitive, behavioral, and dopaminergic abnormalities that resemble schizophrenia. Biol Psychiatry. 2011;69(5):432–41.

35. Kleschevnikov AM, Belichenko PV, Faizi M, Jacobs LF, Htun K, Shamloo M, et al. Deficits in cognition and synaptic plasticity in a mouse model of Down syndrome ameliorated by GABAB receptor antagonists. J Neurosci. 2012;32(27):9217–27.

36. Kehrer C, Maziashvili N, Dugladze T, Gloveli T. Altered Excitatory-Inhibitory Balance in the NMDA-Hypofunction Model of Schizophrenia. Front Mol Neurosci. 2008;1:6.

37. Dichter GS, Felder JN, Bodfish JW. Autism is characterized by dorsal anterior cingulate hyperactivation during social target detection. Soc Cogn Affect Neurosci. 2009;4(3):215–26.

38. Matsuo K, Glahn DC, Peluso MA, Hatch JP, Monkul ES, Najt P, et al. Prefrontal hyperactivation during working memory task in untreated individuals with major depressive disorder. Mol Psychiatry. 2007;12(2):158–66.

39. Fuchs T, Jefferson SJ, Hooper A, Yee PH, Maguire J, Luscher B. Disinhibition of somatostatin-positive GABAergic interneurons results in an anxiolytic and antidepressant-like brain state. Mol Psychiatry. 2016.

40. Gandal MJ, Sisti J, Klook K, Ortinski PI, Leitman V, Liang Y, et al. GABAB-mediated rescue of altered excitatory-inhibitory balance, gamma synchrony and behavioral deficits following constitutive NMDAR-hypofunction. Transl Psychiatry. 2012;2:e142.

41. Hearing M, Kotecki L, Marron Fernandez de Velasco E, Fajardo-Serrano A, Chung HJ, Lujan R, et al. Repeated cocaine weakens GABA(B)-Girk signaling in layer 5/6 pyramidal neurons in the prelimbic cortex. Neuron. 2013;80(1):159–70.

42. Marron Fernandez de Velasco E, Hearing M, Xia Z, Victoria NC, Lujan R, Wickman K. Sex differences in GABA(B)R-GIRK signaling in layer 5/6 pyramidal neurons of the mouse prelimbic cortex. Neuropharmacology. 2015;95:353–60.

43. Nimitvilai S, Lopez MF, Mulholland PJ, Woodward JJ. Ethanol Dependence Abolishes Monoamine and GIRK (Kir3) Channel Inhibition of Orbitofrontal Cortex Excitability. Neuropsychopharmacology. 2017;42(9):1800–12.

44. Victoria NC, Marron Fernandez de Velasco E, Ostrovskaya O, Metzger S, Xia Z, Kotecki L, et al. G Protein-Gated K(+) Channel Ablation in Forebrain Pyramidal Neurons Selectively Impairs Fear Learning. Biol Psychiatry. 2016;80(10):796–806.

45. Luján R, Aguado C. Localization and Targeting of GIRK Channels in Mammalian Central Neurons. International review of neurobiology. 2015;123:161–200.

46. Glaaser IW, Slesinger PA. Structural Insights into GIRK Channel Function. International review of neurobiology. 2015;123:117–60.

47. Mombereau C, Kaupmann K, Froestl W, Sansig G, van der Putten H, Cryan JF. Genetic and pharmacological evidence of a role for GABA(B) receptors in the modulation of anxiety- and antidepressant-like behavior. Neuropsychopharmacology. 2004;29(6):1050–62.

48. Fatemi SH, Folsom TD, Thuras PD. Deficits in GABA(B) receptor system in schizophrenia and mood disorders: a postmortem study. Schizophrenia research. 2011;128(1-3):37–43.

49. Yamada K, Iwayama Y, Toyota T, Ohnishi T, Ohba H, Maekawa M, et al. Association study of the KCNJ3 gene as a susceptibility candidate for schizophrenia in the Chinese population. Hum Genet. 2012;131(3):443–51.

50. Cooper A, Grigoryan G, Guy-David L, Tsoory MM, Chen A, Reuveny E. Trisomy of the G protein-coupled K+ channel gene, Kcnj6, affects reward mechanisms, cognitive functions, and synaptic plasticity in mice. Proc Natl Acad Sci U S A. 2012;109(7):2642–7.

51. Lecca S, Pelosi A, Tchenio A, Moutkine I, Lujan R, Herve D, et al. Rescue of GABAB and GIRK function in the lateral habenula by protein phosphatase 2A inhibition ameliorates depression-like phenotypes in mice. Nat Med. 2016;22(3):254–61.

52. Pravetoni M, Wickman K. Behavioral characterization of mice lacking GIRK/Kir3 channel subunits. Genes Brain Behav. 2008;7(5):523–31.

53. Wydeven N, Marron Fernandez de Velasco E, Du Y, Benneyworth MA, Hearing MC, Fischer RA, et al. Mechanisms underlying the activation of G-protein-gated inwardly rectifying K+ (GIRK) channels by the novel anxiolytic drug, ML297. Proc Natl Acad Sci U S A. 2014;111(29):10755–60.

54. Yao G, Chen XN, Flores-Sarnat L, Barlow GM, Palka G, Moeschler JB, et al. Deletion of chromosome 21 disturbs human brain morphogenesis. Genetics in medicine : official journal of the American College of Medical Genetics. 2006;8(1):1–7.

55. Valetto A, Orsini A, Bertini V, Toschi B, Bonuccelli A, Simi F, et al. Molecular cytogenetic characterization of an interstitial deletion of chromosome 21 (21q22.13q22.3) in a patient with dysmorphic features, intellectual disability and severe generalized epilepsy. European journal of medical genetics. 2012;55(5):362–6.

56. Lazary J, Juhasz G, Anderson IM, Jacob CP, Nguyen TT, Lesch KP, et al. Epistatic interaction of CREB1 and KCNJ6 on rumination and negative emotionality. European neuropsychopharmacology : the journal of the European College of Neuropsychopharmacology. 2011;21(1):63–70.

57. Signorini S, Liao YJ, Duncan SA, Jan LY, Stoffel M. Normal cerebellar development but susceptibility to seizures in mice lacking G protein-coupled, inwardly rectifying K+ channel GIRK2. Proc Natl Acad Sci U S A. 1997;94(3):923–7.

58. Kotecki L, Hearing M, McCall NM, Marron Fernandez de Velasco E, Pravetoni M, Arora D, et al. GIRK Channels Modulate Opioid-Induced Motor Activity in a Cell Type- and Subunit-Dependent Manner. J Neurosci. 2015;35(18):7131–42.

59. Anderson EM, Gomez D, Caccamise A, McPhail D, Hearing M. Chronic unpredictable stress promotes cell-specific plasticity in prefrontal cortex D1 and D2 pyramidal neurons. Neurobiol Stress. 2019;10:100152.

60. Deacon RM, Rawlins JN. T-maze alternation in the rodent. Nature protocols. 2006;1(1):7–12.

61. Brady AM, Floresco SB. Operant procedures for assessing behavioral flexibility in rats. J Vis Exp. 2015(96):e52387.

62. Richardson NR, Roberts DC. Progressive ratio schedules in drug self-administration studies in rats: a method to evaluate reinforcing efficacy. J Neurosci Methods. 1996;66(1):1–11.

63. Kara NZ, Stukalin Y, Einat H. Revisiting the validity of the mouse forced swim test: Systematic review and meta-analysis of the effects of prototypic antidepressants. Neurosci Biobehav Rev. 2018;84:1–11.

64. Castagne V, Moser P, Roux S, Porsolt RD. Rodent models of depression: forced swim and tail suspension behavioral despair tests in rats and mice. Current protocols in pharmacology. 2010;Chapter 5:Unit 5.8.

65. Diamond A. Executive functions. Annu Rev Psychol. 2013;64:135–68.

66. Etkin A, Gyurak A, O’Hara R. A neurobiological approach to the cognitive deficits of psychiatric disorders. Dialogues Clin Neurosci. 2013;15(4):419–29.

67. Bissonette GB, Roesch MR. Neurophysiology of rule switching in the corticostriatal circuit. Neuroscience. 2017;345:64–76.

68. Ragozzino ME, Detrick S, Kesner RP. Involvement of the prelimbic-infralimbic areas of the rodent prefrontal cortex in behavioral flexibility for place and response learning. J Neurosci. 1999;19(11):4585–94.

69. Vertes RP. Differential projections of the infralimbic and prelimbic cortex in the rat. Synapse. 2004;51(1):32–58.

70. Marquis JP, Killcross S, Haddon JE. Inactivation of the prelimbic, but not infralimbic, prefrontal cortex impairs the contextual control of response conflict in rats. Eur J Neurosci. 2007;25(2):559–66.

71. Berg L, Eckardt J, Masseck OA. Enhanced activity of pyramidal neurons in the infralimbic cortex drives anxiety behavior. PLoS One. 2019;14(1):e0210949.

72. Godsil BP, Kiss JP, Spedding M, Jay TM. The hippocampal-prefrontal pathway: the weak link in psychiatric disorders? European neuropsychopharmacology : the journal of the European College of Neuropsychopharmacology. 2013;23(10):1165–81.

73. Ishii-Takahashi A, Takizawa R, Nishimura Y, Kawakubo Y, Kuwabara H, Matsubayashi J, et al. Prefrontal activation during inhibitory control measured by near-infrared spectroscopy for differentiating between autism spectrum disorders and attention deficit hyperactivity disorder in adults. NeuroImage Clinical. 2014;4:53–63.

74. Szczepanski SM, Knight RT. Insights into human behavior from lesions to the prefrontal cortex. Neuron. 2014;83(5):1002–18.

75. Marron Fernandez de Velasco E, Carlblom N, Xia Z, Wickman K. Suppression of inhibitory G protein signaling in forebrain pyramidal neurons triggers plasticity of glutamatergic neurotransmission in the nucleus accumbens core. Neuropharmacology. 2017;117:33–40.

76. Page CE, Coutellier L. Prefrontal excitatory/inhibitory balance in stress and emotional disorders: Evidence for over-inhibition. Neurosci Biobehav Rev. 2019;105:39–51.

77. Monteiro S, Roque S, de Sa-Calcada D, Sousa N, Correia-Neves M, Cerqueira JJ. An efficient chronic unpredictable stress protocol to induce stress-related responses in C57BL/6 mice. Front Psychiatry. 2015;6:6.

78. Llamosas N, Bruzos-Cidon C, Rodriguez JJ, Ugedo L, Torrecilla M. Deletion of GIRK2 Subunit of GIRK Channels Alters the 5-HT1A Receptor-Mediated Signaling and Results in a Depression-Resistant Behavior. Int J Neuropsychopharmacol. 2015;18(11):pyv051.

79. Maaswinkel H, Gispen WH, Spruijt BM. Effects of an electrolytic lesion of the prelimbic area on anxiety-related and cognitive tasks in the rat. Behav Brain Res. 1996;79(1-2):51–9.

80. Stern CA, Do Monte FH, Gazarini L, Carobrez AP, Bertoglio LJ. Activity in prelimbic cortex is required for adjusting the anxiety response level during the elevated plus-maze retest. Neuroscience. 2010; 170(1):214–22.

81. de Kloet ER, Molendijk ML. Coping with the Forced Swim Stressor: Towards Understanding an Adaptive Mechanism. Neural plasticity. 2016;2016:6503162.

82. De Pablo JM, Parra A, Segovia S, Guillamón A. Learned immobility explains the behavior of rats in the forced swimming test. Physiology & behavior. 1989;46(2):229–37.

83. Gourley SL, Lee AS, Howell JL, Pittenger C, Taylor JR. Dissociable regulation of instrumental action within mouse prefrontal cortex. Eur J Neurosci. 2010;32(10):1726–34.

84. Gilmartin MR, Miyawaki H, Helmstetter FJ, Diba K. Prefrontal activity links nonoverlapping events in memory. J Neurosci. 2013;33(26):10910–4.

85. Euston DR, Gruber AJ, McNaughton BL. The role of medial prefrontal cortex in memory and decision making. Neuron. 2012;76(6):1057–70.

86. Aggleton JP, Hunt PR, Rawlins JN. The effects of hippocampal lesions upon spatial and non-spatial tests of working memory. Behav Brain Res. 1986;19(2): 133–46.

87. Yang ST, Shi Y, Wang Q, Peng JY, Li BM. Neuronal representation of working memory in the medial prefrontal cortex of rats. Molecular brain. 2014;7:61.

88. Prusky GT, West PW, Douglas RM. Reduced visual acuity impairs place but not cued learning in the Morris water task. Behav Brain Res. 2000;116(2):135–40.

89. Seamans JK, Lapish CC, Durstewitz D. Comparing the prefrontal cortex of rats and primates: insights from electrophysiology. Neurotoxicity research. 2008;14(2-3):249–62.

90. Lowery-Gionta EG, Crowley NA, Bukalo O, Silverstein S, Holmes A, Kash TL. Chronic stress dysregulates amygdalar output to the prefrontal cortex. Neuropharmacology. 2018;139:68–75.

91. Shansky RM, Morrison JH. Stress-induced dendritic remodeling in the medial prefrontal cortex: effects of circuit, hormones and rest. Brain Res. 2009;1293:108–13.

92. Ridderinkhof KR, Ullsperger M, Crone EA, Nieuwenhuis S. The role of the medial frontal cortex in cognitive control. Science. 2004;306(5695):443–7.

93. Mayberg HS, Liotti M, Brannan SK, McGinnis S, Mahurin RK, Jerabek PA, et al. Reciprocal limbic-cortical function and negative mood: converging PET findings in depression and normal sadness. The American journal of psychiatry. 1999;156(5):675–82.

94. Winstanley CA, Bachtell RK, Theobald DE, Laali S, Green TA, Kumar A, et al. Increased impulsivity during withdrawal from cocaine self-administration: role for DeltaFosB in the orbitofrontal cortex. Cerebral cortex (New York, NY : 1991). 2009;19(2):435–44.

95. Winstanley CA, Green TA, Theobald DE, Renthal W, LaPlant Q, DiLeone RJ, et al. DeltaFosB induction in orbitofrontal cortex potentiates locomotor sensitization despite attenuating the cognitive dysfunction caused by cocaine. Pharmacol Biochem Behav. 2009;93(3):278–84.

96. Liston C, Miller MM, Goldwater DS, Radley JJ, Rocher AB, Hof PR, et al. Stress-induced alterations in prefrontal cortical dendritic morphology predict selective impairments in perceptual attentional set-shifting. J Neurosci. 2006;26(30):7870–4.

97. Anisman H, Matheson K. Stress, depression, and anhedonia: caveats concerning animal models. Neurosci Biobehav Rev. 2005;29(4-5):525–46.

98. Willner P, Mitchell PJ. The validity of animal models of predisposition to depression. Behav Pharmacol. 2002;13(3):169–88.

99. Hodes GE, Epperson CN. Sex Differences in Vulnerability and Resilience to Stress Across the Life Span. Biol Psychiatry. 2019;86(6):421–32.

100. McEwen BS. Hormones and behavior and the integration of brain-body science. Hormones and behavior. 2020;119:104619.

101. Jaggar M, Rea K, Spichak S, Dinan TG, Cryan JF. You’ve got male: Sex and the microbiota-gut-brain axis across the lifespan. Frontiers in neuroendocrinology. 2020;56:100815.

102. Pinares-Garcia P, Stratikopoulos M, Zagato A, Loke H, Lee J. Sex: A Significant Risk Factor for Neurodevelopmental and Neurodegenerative Disorders. Brain sciences. 2018;8(8).

103. Beck KD, Luine VN. Sex differences in behavioral and neurochemical profiles after chronic stress: role of housing conditions. Physiology & behavior. 2002;75(5):661–73.

104. Bowman RE, Beck KD, Luine VN. Chronic stress effects on memory: sex differences in performance and monoaminergic activity. Hormones and behavior. 2003;43(1):48–59.

105. Bowman RE, Micik R, Gautreaux C, Fernandez L, Luine VN. Sex-dependent changes in anxiety, memory, and monoamines following one week of stress. Physiology & behavior. 2009;97(1):21–9.

106. Bowman RE, Zrull MC, Luine VN. Chronic restraint stress enhances radial arm maze performance in female rats. Brain Res. 2001;904(2):279–89.

107. Luine V, Gomez J, Beck K, Bowman R. Sex differences in chronic stress effects on cognition in rodents. Pharmacol Biochem Behav. 2017;152:13–9.

108. McLaughlin KJ, Baran SE, Wright RL, Conrad CD. Chronic stress enhances spatial memory in ovariectomized female rats despite CA3 dendritic retraction: possible involvement of CA1 neurons. Neuroscience. 2005;135(4):1045–54.

109. Conrad CD, Grote KA, Hobbs RJ, Ferayorni A. Sex differences in spatial and non-spatial Y-maze performance after chronic stress. Neurobiol Learn Mem. 2003;79(1):32–40.

110. Kitraki E, Kremmyda O, Youlatos D, Alexis MN, Kittas C. Gender-dependent alterations in corticosteroid receptor status and spatial performance following 21 days of restraint stress. Neuroscience. 2004;125(1):47–55.

111. Moench KM, Breach MR, Wellman CL. Prior stress followed by a novel stress challenge results in sex-specific deficits in behavioral flexibility and changes in gene expression in rat medial prefrontal cortex. Hormones and behavior. 2020;117:104615.

112. Moench KM, Wellman CL. Differential dendritic remodeling in prelimbic cortex of male and female rats during recovery from chronic stress. Neuroscience. 2017;357:145–59.

113. Wei J, Yuen EY, Liu W, Li X, Zhong P, Karatsoreos IN, et al. Estrogen protects against the detrimental effects of repeated stress on glutamatergic transmission and cognition. Mol Psychiatry. 2014;19(5):588–98.

114. Garrett JE, Wellman CL. Chronic stress effects on dendritic morphology in medial prefrontal cortex: sex differences and estrogen dependence. Neuroscience. 2009;162(1):195–207.

115. Yohn CN, Ashamalla SA, Bokka L, Gergues MM, Garino A, Samuels BA. Social instability is an effective chronic stress paradigm for both male and female mice. Neuropharmacology. 2019;160:107780.

116. Snyder K, Barry M, Plona Z, Ho A, Zhang XY, Valentino RJ. The impact of social stress during adolescence or adulthood and coping strategy on cognitive function of female rats. Behav Brain Res. 2015;286:175–83.

117. Urban KR, Valentino RJ. Age- and Sex-Dependent Impact of Repeated Social Stress on Intrinsic and Synaptic Excitability of the Rat Prefrontal Cortex. Cerebral cortex (New York, NY : 1991). 2017;27(1):244–53.

